# Oscillatory and elevated flow distinctly regulate gene expression in human coronary artery endothelial cells

**DOI:** 10.1101/2025.10.10.681055

**Authors:** Giulio Vidotto, Sara Luzzi, Jon Humphries, Robert Beal, David G McVey, Christopher P Nelson, Thomas R Webb, Martin J Humphries, Graham R Smith, Stephen J White

**Affiliations:** Bioinformatics Support Unit, Faculty of Medical Sciences, Newcastle University, Newcastle upon Tyne NE2 4HH, UK; Biosciences Institute, Faculty of Medical Sciences, Newcastle University, Newcastle upon Tyne NE2 4HH, UK; Department of Life Sciences, Manchester Metropolitan University, Manchester, UK M1 5GD; Leicester British Heart Foundation Centre of Research Excellence & Division of Cardiovascular Sciences, University of Leicester, and National Institute for Health Research Leicester Biomedical Research Centre, Leicester, UK; Wellcome Centre for Cell-Matrix Research, Faculty of Biology, Medicine & Health, University of Manchester, Manchester M13 9PT, UK

**Keywords:** Endothelial cells, shear stress, plaque erosion, gene expression

## Abstract

**Background:** Atherosclerosis develops at arterial sites exposed to disturbed flow, while plaque rupture and plaque erosion predominantly occur in regions subjected to elevated flow. The impact of elevated flow on regulation of endothelial gene expression is less well studied; therefore, we undertook a comprehensive analysis of primary human coronary artery endothelial cell (HCAEC) gene expression under elevated flow, comparing it to gene expression induced by normal physiological and oscillatory flow.

**Methods:** Analysis of HCAEC mRNA, microRNA and protein expression cultured under oscillatory shear stress (OSS), physiological laminar shear stress (LSS), and elevated shear stress (ESS) for 72 hours. Identification of changes in RNA isoform expression and proximity of flow-responsive genes to established coronary artery disease (CAD) risk loci were also performed.

**Results:** 2,175 shear-regulated genes were identified, with 665 uniquely responsive to ESS. Both ESS and OSS induced significant changes in RNA isoform selection, predicted to affect 848 and 580 genes respectively. Signalling pathways regulating CAD pathogenesis including HIPPO, TGFβ/BMP, and IRF, showed altered RNA isoform selection which may influence plaque development and plaque erosion. 65% of linkage disequilibrium (LD)-filtered CAD-associated genetic variants contained at least one OSS or ESS-regulated gene within 250Kb. Proteomic analysis identified 289 proteins differentially expressed under OSS and 171 under ESS, with notable discordance between mRNA and protein changes observed in 28.7% (OSS vs LSS) and 16.6% (ESS vs LSS) genes. Additionally, 40 shear-responsive microRNAs were identified.

**Conclusion:** Elevated flow elicits a distinct gene expression programme in HCAECs, modulating pathways central to CAD pathogenesis.

**Research Perspective:** **What New Question Does This Study Raise?**

- Plaque erosion predominantly occurs on the upstream surface of the plaque, where the endothelium is exposed to elevated flow, which we show within this study to evoke a significant and largely unique regulation of the transcriptome, miRome and proteome within primary human coronary artery endothelial cells.
- Identify shear stress as a significant regulator of alternative splicing, with elevated flow causing the greatest shift in alternative transcript selection.
- Identify 135 oscillatory, and 101 elevated differentially expressed genes in proximity to CAD risk loci.

**What Question Should be Addressed Next?**

- Study of the response of the coronary endothelium directly in patients to understand how the risk factors involved in plaque erosion change endothelial function.
- Creation of physiological 3D arterial models replicating in vivo observations to study the precipitating factors involved in plaque erosion.

## Introduction

Coronary artery disease (CAD) and its acute and chronic clinical manifestations remain one of the greatest contributors of worldwide morbidity and mortality. The haemodynamic environment modifies many aspects of endothelial behaviour, with low average shear stress and disturbed ‘athero-prone’ flow promoting plaque development and progression^1–5^. By contrast, plaque disruption most frequently occurs in regions of the plaque exposed to elevated flow. This is true for both plaque rupture and plaque erosion, which represent the most frequent contributors to atherothrombotic causes of myocardial infarction and predominantly occur on the upstream surface and area of minimal lumen area (MLA) of stenotic plaques where the endothelium encounters elevated flow^6–8^.

While the endothelial response to disturbed or oscillatory flow has received significant attention, the response of the endothelium to elevated flow, where ACS most frequently occurs, has not been studied in depth^9^. To address this shortfall, we present an in-depth multi-omic assessment of the response of primary human coronary artery endothelial cells (HCAECs) to elevated flow, with comparison to normal physiological laminar flow and contrast this to the response to oscillatory flow. Compared to normal physiological flow, elevated and oscillatory flow both significantly regulate gene expression, splicing, microRNA (miR) expression and protein expression, although in distinct and largely non-overlapping manner. Additional analysis revealed a significant enrichment of oscillatory or elevated flow-regulated genes within 250kb of genomic loci known to carry risk for development of Coronary Artery Disease (CAD) identified by GWAS^10,11^.

## Methods

### HCAEC culture and generation of RNA and protein samples for analysis

Confluent monolayers of three independent batches of male HCAECs were exposed to oscillatory ±0.5Pa 1Hz (OSS), normal physiological laminar shear stress (1.5Pa LSS) and elevated laminar shear stress (7.5Pa ESS) for 72 hours using our established parallel plate flow chambers as previously described^9^ ^12^ ^13^ ^14^. In brief, passage 3-4 HCAECs were seeded at high density onto gelatin-coated slides and cultured for 72 hours to ensure confluence and allow stable cell-cell junctions to form and extracellular matrix deposition to occur. Subsequently these HCAECs were cultured under flow for 72 hours before lysis for either RNA or protein purification. The ability to run >25 flow chambers at a time allowed the protein and RNA samples for each individual donor to be prepared in a single experiment, allowing direct comparison across the different omic analyses for each HCAEC donor. Each donor RNA or protein sample represents one replicate in each analysis, therefore there was n=3 RNA or protein samples analysed.

### Bioinformatic processing and data availability

Description of RNAseq, miRseq, splicing analysis, proteomics, CAD-associated loci analysis and machine learning models are included in the supplementary information file. A complete description of the bioinformatic methodology, scripts for each step including merging miR databases and machine learning are available at: https://github.com/GiulioVidotto/Influence_of_miRs_in_regulating_the_response_of_endothelial_cells_to_different_flow_environments

The combined RNAseq and miRseq datasets are available at GSE300941. Proteomics raw data are available at (currently in progress, will update).

## Results

### Pathological flow induces distinct gene expression changes in endothelial cells

We performed multi-omic analyses using HCAECs from three independent male donors. The samples for all the omics on an individual donor were collected at the same time, allowing direct comparisons of data generated across the different omic analyses (Figure 1A). RNAseq demonstrated that compared to physiologically laminar flow of 1.5 Pa (LSS), oscillatory flow ±0.5Pa 1Hz (OSS) uniquely regulated 1218 differentially expressed genes (DEGs) >2-fold, padj<0.05, while elevated laminar flow 7.5Pa (ESS) regulated 675 DEGs >2-fold, padj<0.05. 302 DEGs showed >2-fold regulation by both OSS and ESS (padj<0.05, Figure 1B, 1C). Significantly regulated genes are listed in supplementary data tables 1 & 2. Examination of the expression of arterial markers (SEMA3G, NOTCH4, GJA5 -all highly expressed, GJA4, variable between donors), vein markers (ACKR1, VCAM1-both expressed at low levels), capillary markers (CA4, RGCC, HEY1, F2RL3 -moderate to low expression) summarised in^15^, would suggest the HCAECs used in this study generally maintained an arterial-like phenotype.

**Figure 1.**
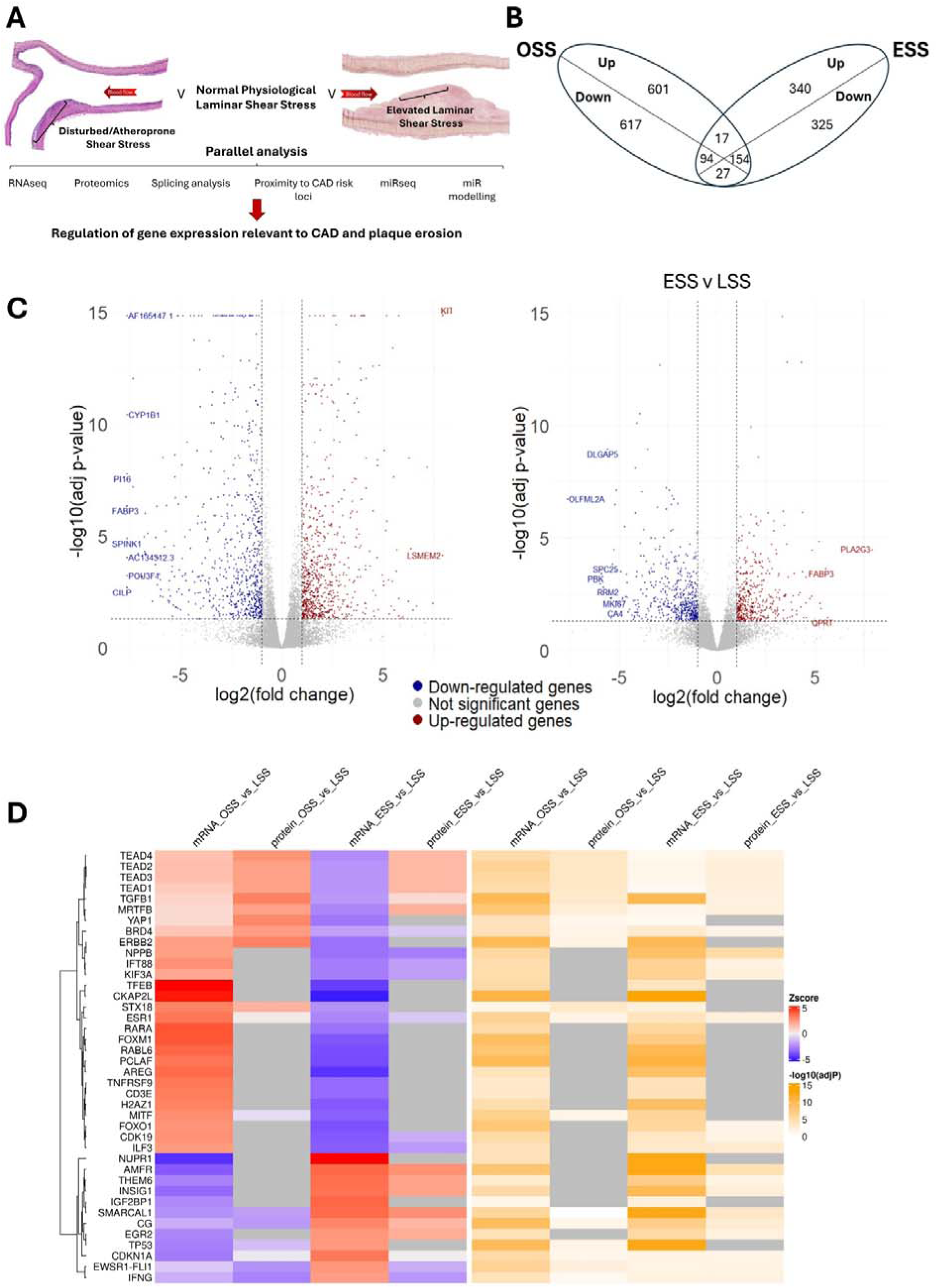
Analysis of the impact of oscillatory and elevated flow on mRNA expression. A. overview of the analysis pipeline. B. Venn diagram of the differentially expressed genes. This plot shows the overlap of differentially expressed genes between Oscillatory Shear Stress (OSS) and Elevated Shear Stress (ESS) conditions. Each section represents up-regulated (Up) or down-regulated (Down) genes that are either unique to one condition or shared between both. C. Volcano plot of the result of differential expression analysis (DEA) for the mRNA-seq in the OSS vs LSS and ESS vs LSS conditions. The x-axis shows the log2 of the fold change (log2FC) values, while the y-axis represents the −log10 of the adjusted p-values. Genes with log2FC ≤ -1 and adjusted p-value ≤ 0.05 are labelled as down-regulated (blue), and genes with log2FC ≥ 1 and adjusted p-value ≤ 0.05 are labelled as up-regulated (red). Genes that do not meet these criteria are considered not significant (grey). The volcano plot for the mRNA-seq in the OSS vs LSS condition is on the left while the one for the mRNA-seq in the ESS vs LSS condition is on the right. D. Heatmap of the top upstream regulators in the OSS vs LSS and ESS vs LSS conditions. This plot shows the top 40 upstream regulators identified using Ingenuity Pathway Analysis (IPA) at the transcriptomics and proteomics levels. The Z-score represents the predicted activation (red) or inhibition (blue) of the upstream regulators, while significance is shown as negative log10 of the adjusted p-values. Grey boxes identify missing data.

A combined analysis of shear regulated genes at the mRNA and protein level (described in detail below) revealed a dominant role for TEADs, YAP, and TGFβ, aligning to previously reported shear regulatory programmes (Figure 1D, genes related to each pathway are listed in supplementary data tables 3 & 4). In addition, Myocardin-related transcription factor B (MRTFB) a co-transcriptional activator of SRF shows a consistent upregulation in OSS and downregulation under ESS. TP53 (p53) and CDKN1A (p21) gene programmes change in a way consistent with the established control of cell cycle by physiological laminar flow.

Of note, there were a number of hypervariable genes, with a >8-fold mean increase or decrease expression in either OSS or ESS that failed to reach significance due to variation between donors (padj>0.05, Supplementary data tables 5 & 6), highlighting genes that constitute a donor-specific response to flow. Further investigation of these hypervariable genes in larger datasets may provide mechanistic insight into patient-specific CAD risk.

### Ingenuity Pathway analysis (IPA)

The IPA pathway analysis provided an interesting insight into the function of OSS and ESS DEGs (Figure 2A & 2B, with pathway-related genes listed in supplementary data file sheets 7 & 8). Repetition within overlapping IPA-identified pathways were removed through simplifying the pathways in R using binary distance between the gene sets associated with each pathway, and hierarchical clustering via the hclust algorithm (Figures S3, S4) with the most significant pathway in each cluster presented in Figure 2. All pathways are listed in supplementary data Figure S5, S6 and networks showing related IPA-identified pathways (Figures S7, S8). Both shear environments were dominated by the regulation of cholesterol synthesis, predicted to be low in OSS and high in ESS. This was consistent with the observed downregulation of INSIG1-dependent gene expression in OSS and upregulation in ESS (Figure 1D). Genes controlling cell cycle are again consistent with the known interaction between laminar flow and cell proliferation.

**Figure 2.**
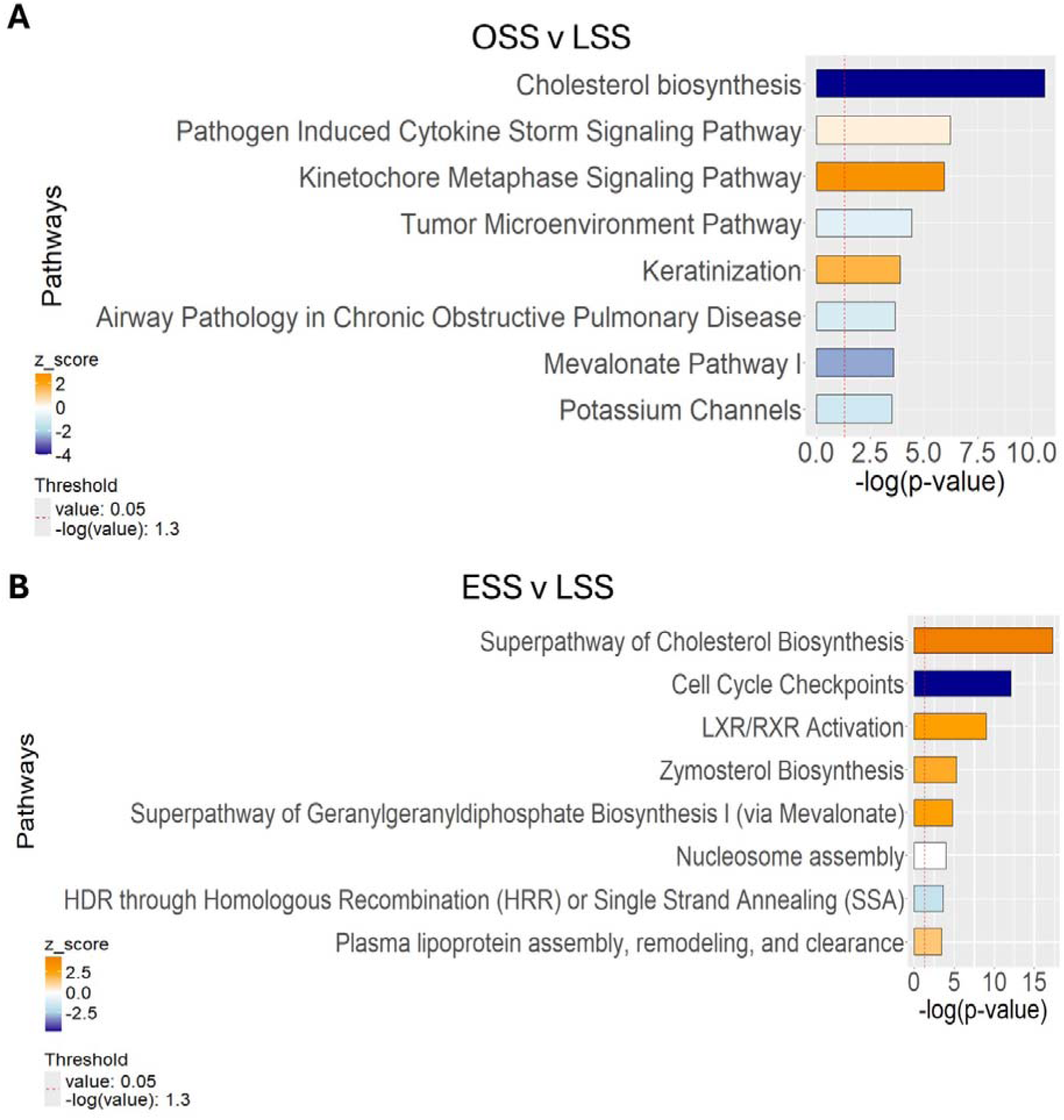
Pathway analysis of differentially regulated genes. A and B. Horizontal bar charts of the canonical pathways at the transcriptomics level in the OSS vs LSS and ESS vs LSS conditions. The plots show the most enriched pathways at the transcriptomics level for the OSS vs LSS (A) and ESS vs LSS (B) biological contrast. On the y-axis are shown the names of the enriched pathways, ordered based on the significance and the x-axis displays the negative log10 of the p-values of overlap. Each bar is colour-coded according to the z-score, with orange and blue bars in the bar chart indicating predicted pathway activation or predicted inhibition, respectively.

### Protein expression and pathway analysis

Liquid chromatography tandem mass spectrometry (LC-MS/MS) analysis of HCAEC proteome identified a total of 597 shear-regulated proteins, compared to 2175 genes by RNAseq (>2-fold, padj<0.05). 120 up-regulated and 169 down-regulated proteins were uniquely regulated between OSS vs LSS, 79 up-regulated and 92 down-regulated proteins were uniquely regulated between ESS vs LSS, while 137 proteins were significantly modulated by both ESS and OSS (Figure 3A-C and supplementary data tables 17 & 18). As with the transcriptomic analysis, there were substantial numbers of distinctly regulated proteins under both OSS and ESS.

**Figure 3.**
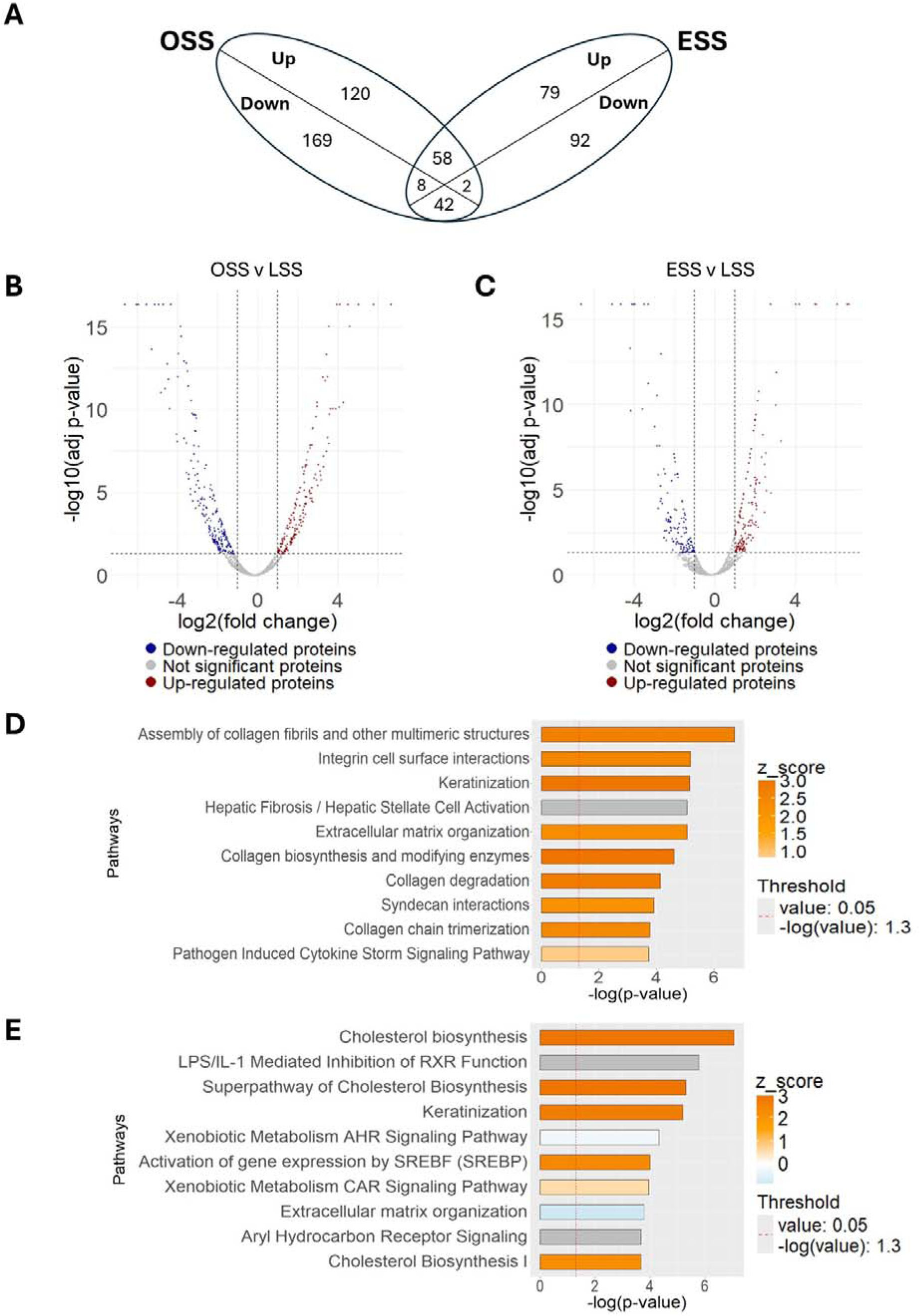
Analysis of the impact of oscillatory and elevated flow on protein expression. A. Venn diagram of the differentially expressed proteins. This plot shows the overlap of differentially expressed proteins between Elevated Shear Stress (ESS) and Oscillatory Shear Stress (OSS) conditions. Each section represents up-regulated (Up) and down-regulated (Down) miRNAs that are either unique to one condition or shared between both. B and C. Volcano plots of the result of differential expression analysis (DEA) for the proteins in the OSS vs LSS and ESS vs LSS conditions. The x-axis shows the log2 of the fold change (log2FC) values, while the y-axis represents the −log10 of the adjusted p-values. Genes with log2FC ≤ -1 and adjusted p-value ≤ 0.05 are labelled as down-regulated (blue), and genes with log2FC ≥ 1 and adjusted p-value ≤ 0.05 are labelled as up-regulated (red). Genes that do not meet these criteria are considered not significant (grey). (B) DEA results for proteomics in the OSS vs LSS contrast. (C) DEA results for proteomics in the ESS vs LSS contrast. D and E. Horizontal bar charts of the canonical pathways at the proteomics level in the OSS vs LSS and ESS vs LSS conditions. On the y-axis are shown the names of the enriched pathways, ordered based on the significance and the x-axis displays the negative log10 of the p-values of overlap. Each bar is colour-coded according to the z-score, with orange and blue bars in the bar chart indicating predicted pathway activation or predicted inhibition, respectively. (D) Canonical pathways identified at the proteomics level for the OSS vs LSS condition. (E) Canonical pathways identified at the proteomics level for the ESS vs LSS condition.

Many pathways related to extracellular matrix organisation and collagen biosynthesis, were significantly regulated at the level of protein expression (Figure 3D, 3E). As this analysis was performed on confluent HCAEC monolayers cultured under flow for 72 hours allowing the accumulation of extracellular matrix, the inclusion of the extracellular matrix within the protein sample provides additional focus on the extracellular compartment allowing the contribution to extracellular matrix to more clearly observed. Consistent with the RNAseq results, cholesterol biosynthesis pathways were activated under ESS. These results add new information in addition to the analysis of gene expression at the RNA level.

### Oscillatory and elevated shear stress have distinct impact on production of alternative RNA isoforms

To further understand the impact of shear stress on gene expression, we sought to gain insights into alternative RNA processing. Alternative RNA splicing plays a key role in cardiovascular function and has been associated to cardiovascular disease including atherosclerosis^16–18^. However, the impact of elevated flow on alternative splicing has not been previously studied. Therefore, we performed differential splicing analyses using rMATS and identified a significant impact of shear stress on regulation of transcript selection. N=3 has sufficient power for this analysis^19^. Compared to LSS, OSS and ESS resulted in 727 or 1158 alternative splicing events respectively (Figure 4A-D). The effect of OSS and ESS on alternative splicing is largely distinct, with only 150 genes (26% of OSS vs LSS and 18% of ESS vs LSS) being differentially spliced in both OSS and ESS, and only 82 splicing events (11% of OSS vs LSS and 7% of ESS vs LSS) appear differentially regulated in both OSS and ESS compared to LSS. The majority of predicted splicing changes involve exon skipping/inclusion, with ESS leading to a particularly high number of exon skipping events however, OSS resulted in markedly reduced intron retention (Figure 4E, 4F; gene lists presented in supplementary data tables 9 & 10). This analysis spotlights that significant modulation of gene expression in response to shear stress occurs through regulation of the abundance of alternative RNA isoforms, in addition to overall transcriptional control. Gene Ontology analyses revealed a number of common themes including regulation of cell surface receptors, cell-cell junction components, membrane and cytoskeletal organisation. Of potential importance for plaque erosion, elevated flow modulated splicing of genes involved in cell substrate adhesion, integrin-mediated signalling, response to reactive oxygen species and TOR signalling (Figure 4G, 4H and supplementary data tables 11 & 12).

**Figure 4.**
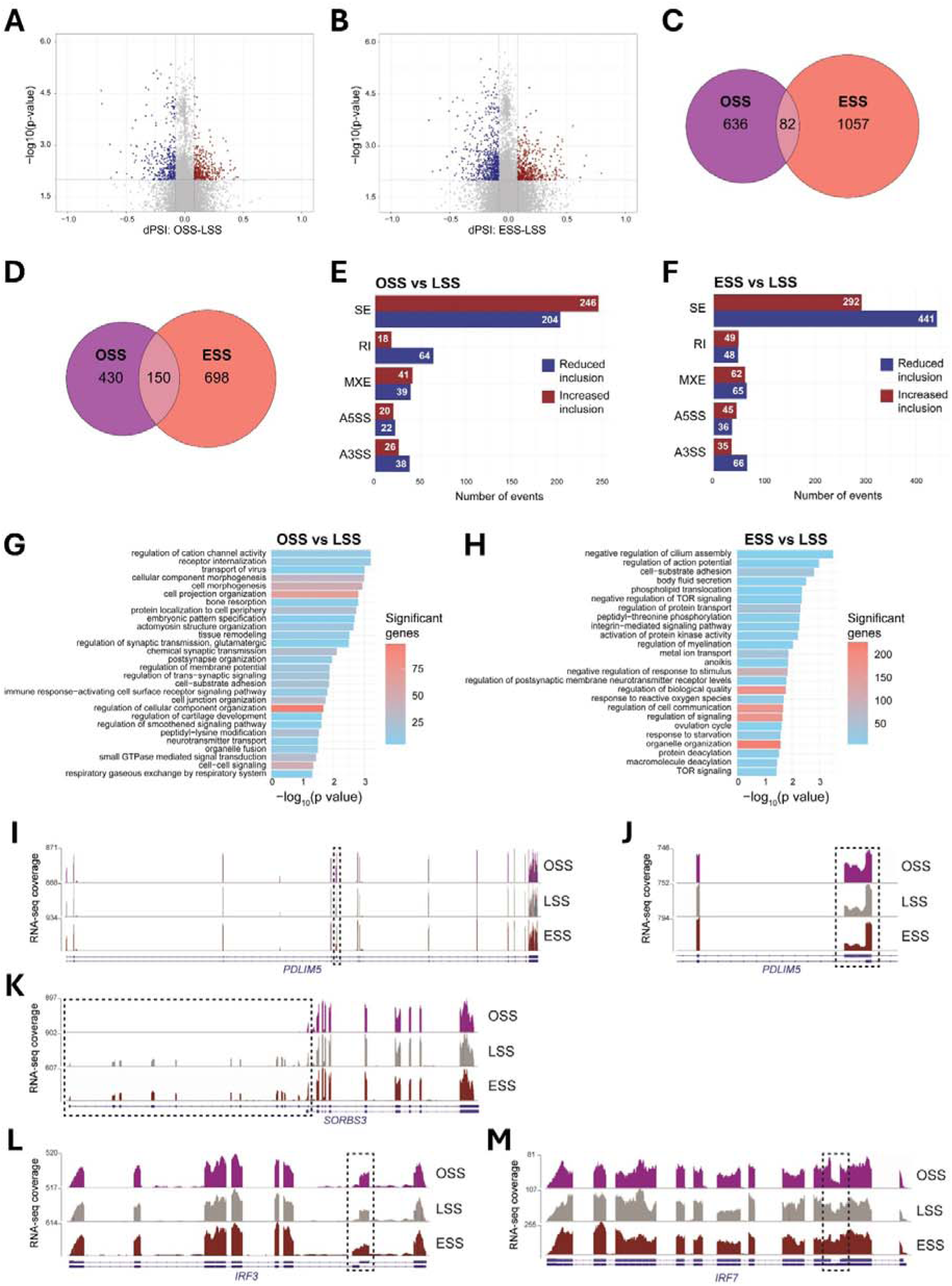
The impact of OSS and ESS on RNA transcript selection. A. Volcano plot shows alternative splicing events within protein-coding genes identified in OSS compared to LSS as calculated using rMATS v.4.1.2. dPSI, differential percentage splicing inclusion. Blue and red dots represent splicing events with significantly reduced (dPSI ≤ -0.08 and p-value < 0.01) or significantly increased inclusion (dPSI ≥ 0.08 and p-value < 0.01) in OSS compared to LSS. B. Analysis as in A, but for alternative splicing events identified in ESS compared to LSS. C, D. Venn diagrams show common differential alternative splicing events (C) or common genes undergoing differential alternative splicing (D) in both OSS and ESS compared to LSS. E. Scatterplot analysis identifies alternative splicing events significantly changing in both OSS and ESS compared to LSS. Red dots indicate genes with opposite effect of OSS and ESS on the same splicing inclusion events. F. Bar plot illustrates types of significant alternative splicing events identified in OSS compared to LSS. For each, the number of events showing significantly reduced (blue bars) and significantly increased (red bars) inclusion levels are indicated. SE, exon skipping/inclusion. RI, intron retention. MXE, mutually exclusive exons. A5SS, alternative 5′ splice site. A3SS, alternative 3′ splice site. G. Analysis as in E, but for alternative splicing events identified in ESS compared to LSS. H. Bar plot shows gene ontology analysis of genes that undergo alternative splicing in OSS compared to LSS. Colours indicate the number of significant genes belonging to the respective biological process. I. Analysis as in G, but for alternative splicing events identified in ESS compared to LSS. L-O. Snapshot of RNA-seq merged tracks from the three batches of HCAECs subjected to OSS, LSS and ESS from the IGV genome browser showing significant differential alternative splicing events over *PDLIM5*, *SORBS3*, *IRF3* and *IRF7*.

Of relevance to CAD, the YAP/TAZ signalling regulators PDLIM5 and SORBS3 displayed OSS-specific alternative splicing patterns (Figure 4I-K). Specifically, OSS but not ESS led to increased inclusion of an alternative 3’ splice sites within *PDLIM5* RNA predicted to introduce 109 amino acids within the disordered domain, compared to the protein isoform expressed in LSS. Furthermore, OSS caused a dramatic switch in *SORBS3* isoform expression leading to a shorter RNA transcript expressing a protein isoform that lacks the SoHo (Sorbin Homology) domain, which enables interactions with cytoskeletal proteins like vinculin^20^. On the other hand, ESS but not OSS also caused significant changes in alternative splicing patterns of YAP/TAZ activators, in particular *TEAD1* and *PARD3* (supplementary data tables 9 & 10), associated to a shorter TEAD1 protein isoform and a reduced hinge between the PARD3 PDZ domains. Interestingly, ESS and OSS also led to changes in the interferon regulators *IRF3* and *IRF7* respectively (Fig 4L, 4M), both affecting selection of the first methionine during protein translation. ESS caused increased selection of a mutually exclusive exon with predicted production of a shorter IRF3 protein isoform lacking the N-terminal domain, which includes the Nuclear Localisation Signal. OSS enhanced recognition of an exitron (internal region of an exon spliced like an intron), which would result in selection of a different N-terminal domain. Production of these alternative isoforms, alongside observed changes in IRF regulators IRAK4 and STING1 (supplementary data tables 9 & 10), may alter sensitivity of IRF signalling, implicated to play a role in plaque erosion^21^.

### Expression and regulation of putative CAD risk loci genes in HCAEC

GWAS has identified numerous genetic loci that associate with variable risk of CAD. To determine whether the differentially-expressed genes (DEGs) identified in the OSS and ESS conditions were located near to CAD risk loci, a list of linkage disequilibrium (LD)-filtered variants from two recent CAD multi-population meta-analyses was compiled ^10,11^ (375 variants in total with MAF>0.005) and calculated the distance between DEG start sites and CAD variants. 1495 OSS DEGs and 954 ESS DEGs from chr 1-22 and chr X were considered in this analysis. 135 OSS DEGs and 101 ESS DEGs were located within 250kb of a CAD risk variant. 36% of the CAD-associated variants had at least one OSS DEG within 250kb, whilst 26.93% were within 250kb of at least one of the ESS DEGs. A summary of the DEGs within 250kb of the CAD variants is shown in figure 5A, genes and SNPs listed in supplementary data tables 13 & 14. Circos plots indicating the approximate locations of all OSS and ESS DEGs within 250kb of CAD-associated variants are shown in Figures 4B,C respectively.

**Figure 5.**
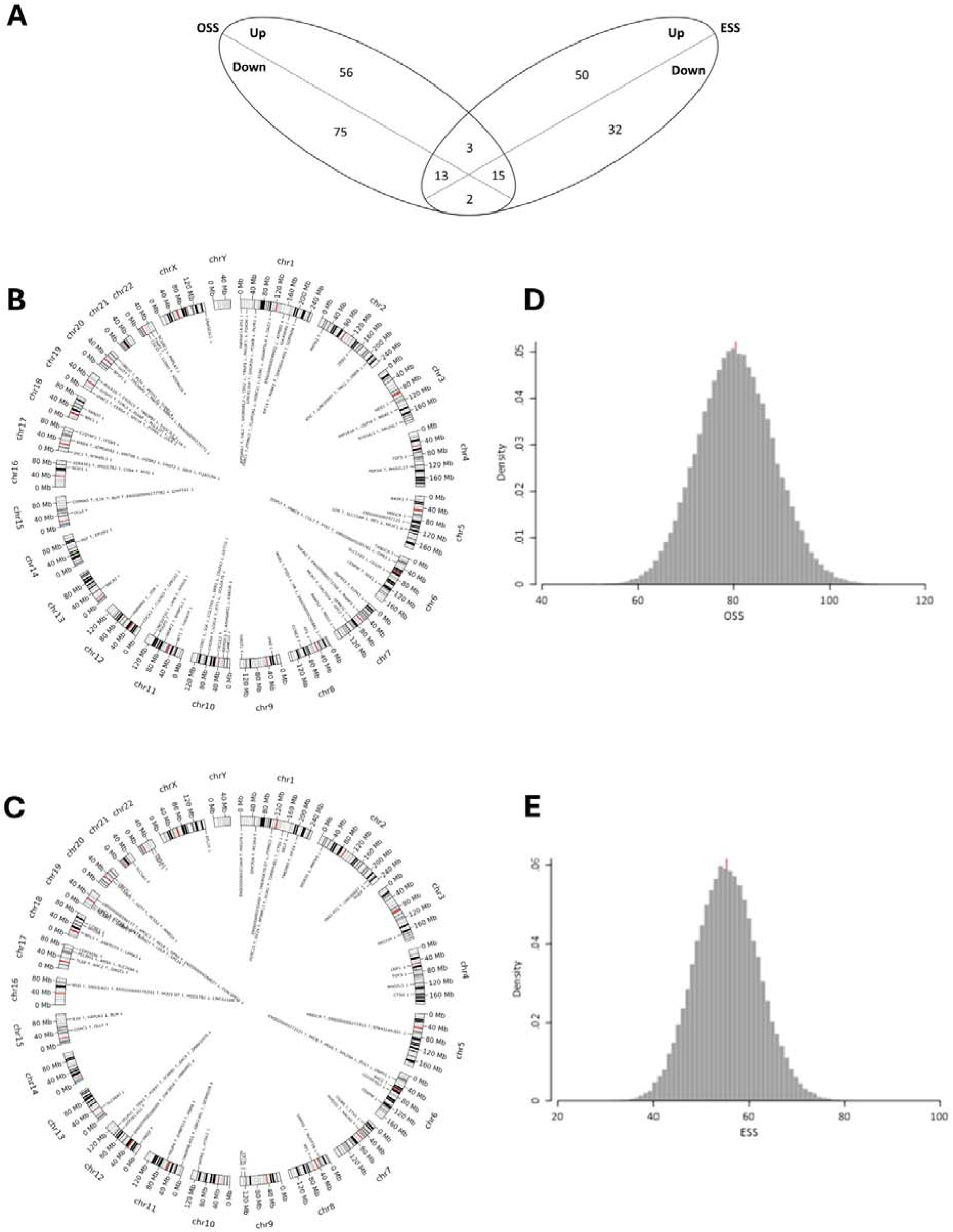
Proximity of OSS and ESS differentially regulated genes to CAD risk loci. A. Venn diagram of the OSS and ESS differentially-expressed genes within 250kb of a CAD-associated variant. This plot shows the number of differentially-expressed genes under OSS and ESS conditions whose gene start site is located within 250kb of a CAD-associated variant. Each section indicates the shear stress type (OSS or ESS) and the direction of the gene expression change under the indicated condition compared to LSS. B. Circos plot showing the approximate location of OSS differentially-expressed genes within 250kb of a CAD-associated variant. Circos plots showing the approximate locations of all OSS differentially-expressed genes (compared to laminar flow) within 250kb of a CAD-associated variants. Up arrows indicate genes whose expression was increased under OSS vs LSS, whilst down arrows indicate genes whose expression was decreased under OSS vs LSS. Locations are approximate to ensure legibility of all labels. C. Circos plot showing the approximate location of ESS differentially-expressed genes within 250kb of a CAD-associated variant. Circos plots showing the approximate locations of all ESS differentially-expressed genes (compared to laminar flow) within 250kb of a CAD-associated variants. Up arrows indicate genes whose expression was increased under ESS vs LSS, whilst down arrows indicate genes whose expression was decreased under ESS vs LSS. Locations are approximate to ensure legibility of all labels. D. Histogram showing the permutation distribution of OSS. The results from 100,000 permutations identifying differentially-expressed genes (compared to laminar flow) near randomly selected genetic variants. The mean is shown in red with the maximum for OSS random variants being 114, compared with 135 found for CAD variants. E. Histogram showing the permutation distribution of ESS. The results from 100,000 permutations identifying differentially-expressed genes (compared to laminar flow) near randomly selected genetic variants. The mean is shown in red with the maximum for ESS random variants being 87, compared to 101 found for CAD variants.

An enrichment analysis was performed to investigate whether the association between the 375 CAD lead SNPs and shear-regulated DEGs might occur by chance. Repeated analysis of 375 similar, but randomly selected SNPs 100,000 times identified the probability of observing the association with CAD risk loci by random chance was extremely small (p<1x10^-6^ for both OSS and ESS, Figures 5D, E). This highlights a potential role for shear regulated DEGs in participating in the genetically identified pathogenesis of CAD.

### The haemodynamic environment modifies the expression of miRs

Sequencing of the miRome of HCAECs identified that HCAECs express a total of 1113 miR. In contrast to changes in mRNA expression, comparatively few miRs were regulated by shear stress (>2-fold, padj<0.05, >10 normalised average counts summed across all conditions, Figure 6A-C) listed Supplementary data tables 15 & 16, again showing limited overlap between the OSS and ESS-regulated miRs. OSS v LSS had the greatest effect on miR expression, regulating a total of 32 miR, with ESS v LSS regulating only 11 miR >2-fold. Despite the relatively small number of significantly regulated miRs, their predicted regulation of genes showing discordance between their mRNA and protein expression was deemed significant (see below) indicating an important role in regulating endothelial gene expression in response to shear stress.

**Figure 6.**
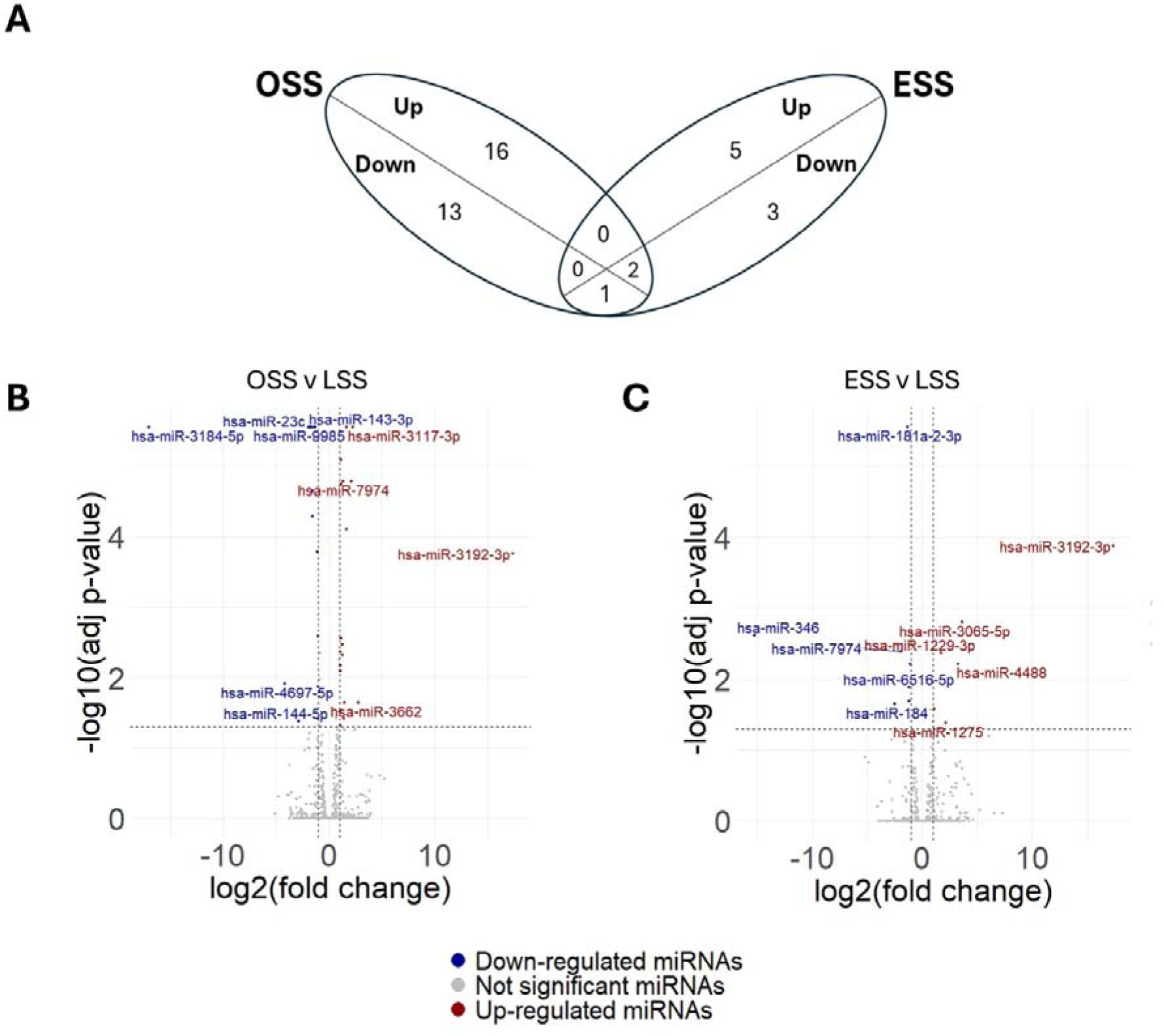
Analysis of the impact of oscillatory and elevated flow on miRNA expression. A. Venn diagram of the differentially expressed miRNAs. This plot shows the overlap of differentially expressed miRNAs between Elevated Shear Stress (ESS) and Oscillatory Shear Stress (OSS) conditions. Each section represents up-regulated (Up) and down-regulated (Down) miRNAs that are either unique to one condition or shared between both. B and C. Volcano plots of the result of differential expression analysis (DEA) for the miRNA-seq in the OSS vs LSS and ESS vs LSS conditions. The x-axis shows the log2 of the fold change (log2FC) values, while the y-axis represents the −log10 of the adjusted p-values. Genes with log2FC ≤ -1 and adjusted p-value ≤ 0.05 are labelled as down-regulated (blue), and genes with log2FC ≥ 1 and adjusted p-value ≤ 0.05 are labelled as up-regulated (red). Genes that do not meet these criteria are considered not significant (grey). (B) DEA results for miRNAs in the OSS vs LSS contrast. (C) DEA results for miRNAs in the ESS vs LSS contrast. D. Tables of the up- and down-regulated miRNAs in the OSS vs LSS and ESS vs LSS conditions. Each table has three columns: the first column is about the IDs of the miRNAs; the second one is about the log2(FC) and the last one is about the adjusted p-values associated to each miRNA.

### Comparison between mRNA and protein expression identified genes with discordant gene expression

We aligned the expression of significantly detected proteins with their cognate mRNAs to examine concordance between mRNA and protein expression. 2122 genes (49.9 % of all detected genes) were not regulated at either mRNA or protein expression by any flow environment. 145 flow regulated genes demonstrated a concordant change in both mRNA and protein expression by shear stress. By contrast, 28.7% (OSS vs LSS) and 16.6% (ESS vs LSS) of flow-regulated genes demonstrate discordant regulation between RNA and protein expression (listed in supplementary data tables 19-24, supplementary data Figure S9-S10, top 5 genes in each group presented in Table S1).

To examine whether targeted protein degradation was potentially responsible for discordant gene expression, we examined if these discordantly regulated genes were enriched for KFERQ-like motifs using KFERQ finder V8.0 (https://rshine.einsteinmed.edu/). No association between KFERQ-like motif-containing proteins and discordant gene regulation was observed (supplementary data Figure S11), providing no evidence that alteration of chaperone mediated autophagy is responsible for discordant mRNA-protein expression.

### Assessment of the impact of miRs in the regulation of discordant gene expression

A list of predicted gene targets for the shear-regulated miRs was generated by combining miRDB v6.016, TarBase v9.017, MirTarBase v9.018 and TargetScan v8.0 databases. Subsequent analysis of enrichment of these predicted gene targets with the discordantly regulated genes failed to find any association (Supplementary Figures S11), suggesting that potential miR-gene interaction alone is insufficient to identify any targeting of the discordant gene expression and that other factors influence their interaction.

Exploratory analysis using multiple computational models identified that a random forest classifier was able to identify an interaction between shear regulated miRs and discordantly regulated gene expression, with AUC scores of 0.71, for both OSS and ESS subsets of genes (Figure 7A-B). This required the inclusion of additional data, including the relative read counts of target mRNAs, the number of shear regulated miRs that were predicted to target an individual gene, the total read counts of the targeting miRs and the calculated binding energy of the miR-mRNA interaction. SHAP values were calculated and visualised to interpret feature contributions to the prediction of the Random Forest model, highlighting that the mRNA read count (equivalent to the concentration) had the greatest contribution to the classifier (Figure 7C-D). While difficult to predict on an individual miR-gene interaction, this analysis does support an important role for shear-regulated miRs in controlling the expression of genes where there is discordance between the mRNA and protein abundance.

**Figure 7.**
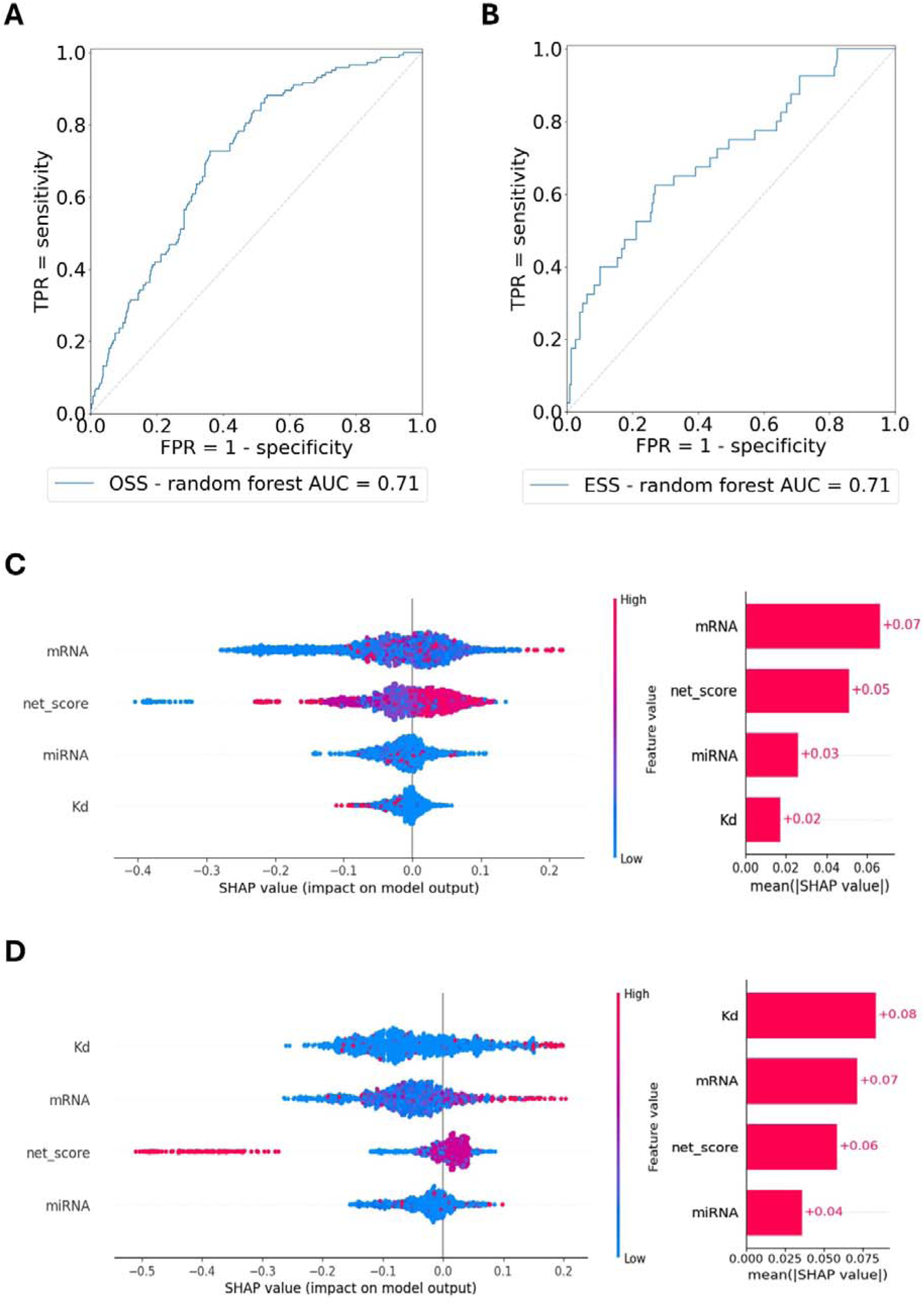
Machine learning models to investigate the impact of shear-regulated miRNAs on discordant gene regulation. A and B. ROC curve on the OSS and ESS samples. The plots show the performances of the random forest models on the test set. The x-axis shows the false positive rate (FPR) given by subtracting all the specificity values from one. The y-axis shows the true positive rate (TPR) calculated from the sensitivity. (A) Performance of the random forest classifier on the samples exposed to OSS. (B) Performance of the random forest classifier on the samples exposed to ESS. C and D. SHAP values beeswarm plots and bar plots. On the right, the beeswarm plots display an information-dense summary of how the top features in a dataset impact the model’s output. A single dot on each feature row represents each SHAP value related to a feature. The SHAP value of that feature determines the x position of the dot, and dots “pile up” along each feature row to show density. Colour is used to display the original value of a feature. (C) SHAP value distribution of each value of the random forest trained on the OSS samples. (D) SHAP value distribution of each value of the random forest trained on the ESS samples.

## Discussion

The causal link between disturbed flow and both atherogenesis and plaque progression has resulted in a significant body of literature describing how the changes in endothelial gene expression and cell function promote pathology ^5,22^. By contrast, the impact of elevated flow on changes in endothelial gene expression has received less attention to date ^9,12,21^. This study addresses this knowledge gap by providing an in-depth multi-omic analysis of the response of HCAECs to elevated flow, with a full comparison to the better described comparisons between oscillatory and normal physiological flow. Culture of HCAECs under elevated flow alters mRNA, miR and protein expression, with more than half of the regulated genes being uniquely altered under elevated flow compared to either normal physiological laminar flow or oscillatory flow.

In an undiseased artery, elevated flow drives endothelial- and NO-dependent expansive remodelling of the artery^23,24^, however, on the surface of stenotic atherosclerotic plaques, the endothelium is chronically exposed to elevated flow alongside proinflammatory signals, sustaining an altered shift in gene expression. This is potentially important in plaque erosion, which is precipitated by loss of endothelial integrity overlying stenotic atherosclerotic plaques. Most plaque erosions occur on the upstream surface of the plaque and through the area of maximal stenosis, where the endothelium is exposed to elevated flow^6–8,25–27^. Importantly, there doesn’t appear to be a threshold shear stress value above which plaque erosion occurs^6^. Large differences in magnitude of shear stress exist across an individual eroded plaque and also between eroded plaques from different patients, suggesting that elevated flow creates a permissive change in phenotype upon which other factors tigger detachment.

Data presented here in conjunction with our previously published data provide a mechanism through which the response to elevated flow provides a permissive change in cell behaviour that may promote plaque erosion. We have previously shown that elevated flow amplifies the stress response to simulated smoking that is mediated through Nrf2. The resulting shear-dependent upregulation of OSGIN1&2 coordinated a combined Nrf2, HSF1 and IRF-like response, resulting in disrupted proteostasis and loss of adhesion^21^. Data presented here highlights an enhanced IFN (gamma)-like transcriptional response (Figure 1D) under elevated flow potentially contributing to an erosion-permissive gene expression pattern under elevated flow. In addition, elevated flow upregulation of PLA2G3 (Figure 1C) and PLA2G4C, with PTGS2 and PTGDS upregulated by LSS and ESS compared to OSS (supplementary data table 2), provides greater capacity to generate 15dPGJ2, a known activator of both Nrf2 and HSF1^29,30^, again potentially providing an erosion-permissive priming of the cell stress response, while inhibiting inflammasome activation^31^.

Analysis of the whole cell proteome revealed an increase in the hyaluronan receptor CD44 under elevated flow. The underlying matrix of plaques that erode are rich in hyaluronan ^26,28,32,33^, priming endothelial inflammation through activation of both TLR4 and CD44. Both downstream signalling pathways converge, with TLR4-MYD88-IRF activation being augmented by CD44-ERK-AKT signalling, which also increases IRF3 activation^34,35^, synergising with OSGIN-coordinated cell-stress signalling^21^. Concomitantly, elevated flow also increases the expression of *IRF7* (supplementary data table 2), increasing potential for IRF-signalling.

In addition to overall regulation of mRNA expression, both OSS and to a greater extent ESS, caused a shift in transcript isoform selection, extending previous descriptions of flow-induced changes in RNA splicing^18,36^. Strikingly, our analysis identified novel shear-regulated RNA isoforms in human endothelial cells. Of potential relevance to plaque erosion, altered splicing of *FYN*, *NRF1*, *IRF7*, *IRAK4*, *IRF3*, and *STING1* may modify the cell stress response under elevated flow. Similarly, differential splicing of *PXN* and *ITGB4* have the capacity to modify adhesome function. These require further analysis to define the impact on cellular processes relevant for plaque erosion.

Several components of the YAP/TAZ-TEAD (HIPPO) signalling pathway also showed alternative transcript selection under shear stress. The exclusion of the SoHo domain of SORBS3 under OSS, combined with notable expansion of the disordered domain of PDLIM5 may provide additional regulation for the observed increase of YAP/TAZ signalling under disturbed flow^37,38^ shown to increase the development of atherosclerosis in hyperlipidaemic models^39,40^. The implication of the observed regulation of TEAD1 and PARD3 under ESS on this pathway are as yet undetermined.

SMAD1 upregulation under oscillatory flow increases the capacity for TGFβ/BMP signalling. Similarly, a decrease in CD109 mRNA and protein expression under oscillatory flow would be predicted to promote TGFβ signalling through reducing TFGβ receptor internalisation^41^. Both observations are consistent with disturbed flow providing a more permissive environment of endothelial-to-mesenchymal transition (EndoMT) to occur^42–44^. Mild alteration of splicing observed for both ACVRL1 (OSS) and ACVR1 (ESS) under have the capacity to provide an additional layer of shear modulation of this important pathway that regulates EC function^45,46^ ^47^.

Other notable changes in protein expression included several regulators of coagulation, with downregulation Tissue Factor Pathway Inhibitor (TFPI) and urokinase-type plasminogen activator PLAU under ESS which may promote a more coagulation-permissive environment. ADAMTS1 mRNA and protein downregulation under OSS may potentiate VEGF signalling and angiogenic potential of ECs in this flow environment. Lastly, Medin precursor protein MFGE8 was upregulated under ESS.

Our analysis highlights significant enrichment of both OSS and ESS DEGs at genetic loci associated with CAD risk. Of the 375 LD-filtered variants with MAF>0.005 from two recent CAD multi-population meta-analyses^10,11^, 36% of the CAD-associated variants had at least one OSS DEG within 250kb, whilst 26.9% were within 250kb of at least one of the ESS DEGs. Overall, 246 (65%) loci had at least one OSS or ESS DEG within the same LD block and within 250kb of the lead SNP. The mechanism through which these CAD-associated variants mediate their effects on the pathogenesis of atherosclerosis is largely unknown, however this analysis provides a rationale for further study in HCAECs to determine any regulation of the known athero-protective effects of laminar flow or plaque progression in regions exposed to elevated flow.

The increase in Cholesterol Biosynthesis under LSS and amplified in ESS may denote a requirement enhanced membrane rigidity to withstand higher uni-directional forces, or to ‘desensitise’ mechanosensitive cell surface receptors upon achieving a homeostatic response by increasing membrane stiffness. The reciprocal decrease in Cholesterol Biosynthesis under OSS may promote membrane fluidity, allowing greater sensitivity of mechanosensors. The necessity for synthesis of cholesterol rather than simple uptake and utilisation from LDL is unclear and may point to intermediate metabolites playing an important role in endothelial function, implicated by the pleiotropic effects of statins on endothelial cells^48,49^. In addition, recent identification of cholesterol-related genetic variants with impact on endothelial function were identified^50^. The resultant polygenic risk score highlighted individuals who derived greater benefit from aggressive statin treatment, again stressing the importance of this axis in regulating endothelial function and resultant impact on CAD.

Using multiple omic c collected at the same time has allowed an exploratory analysis of the impact of shear-regulated miRs to be assessed. The best model performance was obtained using a random forest classifier and required the inclusion of mRNA read counts, the number of shear regulated miRs that were predicted to target an individual gene, the total read counts of the targeting miRs and the calculated binding energy of the miR-mRNA interaction to perform well, suggesting these features are functionally important in determining whether gene expression is affected by miRs. Indeed, one of the biggest contributions to the model was the relative abundance of the target gene mRNA and miR, features that are not currently considered within online target prediction tools. The AUC score of 0.71 suggest that shear regulated miRs do play a role in the expression of genes whose mRNA and protein levels are discordantly regulated by shear stress. Although our classifier achieved good performance, incorporating additional relevant features in the future could further enhance its accuracy and further improvements of the model are needed for a priori predictions whether an individual miR-mRNA interaction will affect gene expression.

This current study has a number of limitations. Firstly, the study was performed with HCAECs from three independent male donors. A lack of high quality HCAECs from female donors has hampered the analysis of sex-specific shear regulation of gene expression. Completion of a sex-specific analysis an immediate aim as it may highlight differently regulated pathways of relevance to plaque erosion. Both the mechanism of regulation and the biological significance of alternative splicing are yet to be elucidated. Significant additional investigation will be necessary to assess the impact of this largely unexplored component of gene regulation in endothelial function and pathophysiology. While a number of OSS and ESS DEGs were identified to locate within 250kb and reside within the same LD block as SNPs associated with CAD risk, further analysis is required to identify if the shear-regulated DEGs represent the causal gene for the association with CAD. The sensitivity of detection of proteins expression by LC-MS/MS is lower for the proteomic analysis compared to the RNAseq analysis, highlighted by an observable relationship between fold change and p value (Figure 6B, C). This limited the identification of genes with discordant mRNA-protein regulation to the genes where the proteins were significantly detected by LC-MS/MS.

In conclusion, the present study extends many previous investigations showing that the flow environment significantly regulates HCAEC gene expression by providing a unique integrated dataset of linked mRNA, miR and proteomic data from 3 donors under both oscillatory, normal physiological and elevated flow. This clearly demonstrates the uniqueness of the HCAEC response to elevated flow at every level of gene expression control. In particular, the impact of altered transcript selection is underexplored and predicted by these data to provide a significant additional layer of gene expression control requiring future investigation. Alongside this, the identification of flow-responsive genes in close proximity to CAD risk loci provide a focus for future investigation to identify the mechanism these loci alter CAD risk. Lastly, we provide evidence that miRs play a significant, but poorly defined role in regulating the expression of genes whose protein expression is discordant from its mRNA. This project highlighted the need for integration of multi-omic analyses to provide a full picture of the response of ECs to shear stress and serves as a resource for the analysis of HCAEC gene regulation by flow.

## Acknowledgments

The authors wish to convey their gratitude to Rachel Scholey at the Bioinformatics Core Facility, University of Manchester, for support with data processing and the support of D. Knight, S. Warwood, E. Keevill, M. Kidd and J.N. Selley at the Bio-MS mass spectrometry core facility (RRID: SCR_020987) in the Faculty of Biology, Medicine and Health at the University of Manchester is gratefully acknowledged.

## Authors contributions

SJW, MJH, JH Conceived and designed the research; RB, SL, JH, DGM, TW Acquired the data and provided key resources for analysis; GV, SL, DGM, CPL, TRW & JH, Performed statistical analysis; SJW, GV, GS, TRW, JH Drafted the manuscript; GV, SL, JH, RB, DGM, CPN, TRW, MJH, GS, SJW Made critical revision of the manuscript for key intellectual content.

