## Additional methods and data for "Oscillatory and elevated flow distinctly regulate gene expression in human coronary artery endothelial cells"

**Multi-omic analysis reveals distinct endothelial gene expression profiles induced by pathological flow**

Supplementary methods

*RNAseq, miRseq data generation and analysis*

TruSeq Stranded RNA 76 bp paired-end sequencing was performed on Illumina HiSeq4000. Sequencing quality was assessed using FastQC (v 0.11.3), FastQ Screen (v 0.9.2), trimmed with Trimmomatic (v 0.36) and mapped against the reference genome (hg38) using STAR (version 2.7.2b). Counts per gene were calculated using STAR annotation from GENCODE 32. Normalisation and differential gene expression tests were performed in DESeq2 (v1.42.1) with confirmation of normalisation through p-value analysis. Transcripts with low counts were removed using independent filtering within DESeq2. The mRNAs with low counts across all samples were automatically excluded by setting the independent Filtering parameter to TRUE.

MicroRNA (miR) library preparation was performed using NEBNext® Multiplex Small RNA Library Prep Set and 76 bp paired-end sequenced performed on Illumina HiSeq4000. The R1 single-end RNA-seq reads were quality assessed using FastQC (v 0.11.3), FastQ Screen (v 0.9.2), quality trimmed and filtered for length with Cutadapt v1.18 (min length 18nt, max length 26nt), and mapped against the reference genome (hg38) using Bowtie (v 1.1.1). Counts per gene were calculated using feature Counts using annotation from miRBasev22. Normalisation, principal component analysis (PCA) and differential gene expression tests were performed in DESeq2 v1.20.0. Among the altered miRNAs, only those with at least 15 reads across all the samples were included within the analysis.

*Alternative splicing analyses*

Differential alternative splicing analyses were performed comparing either oscillatory shear stress (OSS) or elevated shear stress (ESS) to laminar shear stress (LSS) using rMATS v.4.1.2^15^. Alternative splicing events were detected from BAM files aligned by hisat2 v.2.2.1^16^ to the GRCh38 human genome reference (Ensembl v.112) and sorted by coordinates using samtools v.1.5^17^. rMATS analyses were carried out using the paired statistical model with a cutoff (--cstat) of 0.05. Results calculated by counting both the junction and the exon reads (JCEC) were filtered to only include differential splicing events with a p-value ≤ 0.01 and predicted differential PSI (dPSI) of 8%.

Gene Ontology analyses of genes displaying these alternative splicing events were performed using GOstats v.2.60.0^18^ to identified enriched biological processes (GOBP). Enrichment was calculated against genes with an average of at least 5 transcripts per million (TPM) across samples (universe), and p-values were adjusted by hypergeometric test. Redundant terms were filtered using Revigo v.1.8.1^19^ with SimRel semantic similarity measure to only include GOBP with a dispensability score lower than 0.3 and containing between 5 and 500 genes.

All plots were produced in R v.4.1.2 using ggplot2 v3.5.1 and eulerr 7.0.2^20^. Snapshots from the Integrative Genomics Viewer (IGV) browser were generated using representative BAM files merged from the three batches of HCAECs with samtools v.1.5.

*Identification of Differentially-Expressed Genes Within and Near To CAD-Associated Loci*

CAD risk-associated variants were identified from two recent multi-population meta-analyses^10, 11^. Variants were filtered by LD using LDlink’s SNPclip function using the European population and a R^2^ threshold of 0.7. Where LD filtering would be applied to multiple variants, the variant with the lowest p-value in the two studies was selected as the lead variant. Variants not within the LDlink database were included in the final list of lead variants. The gene start site for the differentially-expressed genes was identified using Ensembl’s BioMart function and genes with start sites within 250kb of the CAD-associated variants were identified. Circos plots of differentially-expressed genes near to CAD risk variants were plotted using the Python package pyCirclize (<https://pypi.org/project/pyCirclize/>). Where necessary to ensure legibility, gene labels were grouped together and the direction of the differential-expression is indicated by arrows for each gene.

A permutation test was performed to identify whether there was significant enrichment of CAD variants within 250kb of the start sites of differentially expressed genes. For each of the 100,000 permutations, a random variant was selected from well imputed variants (r^2^>0.3) on the same chromosome for each of the 375 CAD variants (MAF>0.5%) and the number of differentially expressed genes within 250kb of each random variant was counted and the total found. Significance was estimated based on the number of times a permutation total exceeded the observed total.

*Proteomic analysis*

Sample preparation and analysis were performed as previously described^21^. In brief, protein samples (500 μg) obtained in SDS lysis buffer (2% SDS, 50 mM Tris pH 6.8, 10% glycerol) from different flow conditions were sonicated (300 seconds, 30W, Covaris LE220-PLUS), reduced (15 mM DTT, 56 °C, 45 minutes) and alkylated (50 mM iodoacetamide, room temperature, 45 minutes). Proteins were precipitated (4x volume acetone, -20 °C, 16 hours), collected by centrifugation (16,000xg 4 °C, 20 minutes) and resuspended in 0.1% (w/v) Rapigest (Waters) in 100 mM ammonium bicarbonate. Proteins were digested with trypsin at a 1:50 (w/v) ratio (37 °C, 16 hours with shaking at 250 rpm) and then acidified with 1% (v/v) trifluoracetic acid (TFA; 37°C ,45 minutes). Samples were desalted using Oasis HLB cartridges (Waters) according to manufacturer’s instructions, eluting in binding solution: 80 % (v/v) acetonitrile (ACN), 5 % (v/v) TFA, 1M glycolic acid and stored at -20 °C.

For mass spectrometry, peptides were resuspended in 5% (v/v) ACN / 1% (v/v) formic acid and analysed by liquid chromatography (LC) tandem mass spectrometry (LC‐MS/MS) using an UltiMate 3000 Rapid Separation LC (RSLC, Dionex Corporation) coupled to a Q Exactive HF (Thermo Fisher Scientific) mass spectrometer. Peptides were separated using a 75 mm x 250 μm i.d. 1.7 μM CSH C18, analytical column (Waters) with the LC gradient from 95% buffer A (0.1% (v/v) FA in water) and 5% buffer B (0.1% (v/v) FA in acetonitrile) to 7% buffer B at 1 minute, 18% buffer B at 58 minutes, 27% buffer B in 72 minutes and 60% buffer B at 74 minutes at 300 nL min‐1. Peptides were selected for fragmentation automatically by data dependant analysis.

Relative protein abundances were calculated based on ion intensity as implemented in Proteome Discoverer (v2.5; ThermoFisher Scientific) [4], utilising the Protein FDR validator node set at high confidence equating to a 1% FDR cut off. RAW files were searched using SEQUEST-HT against the SwissProt and TREMBL human databases (release-2018_01). Carbamidomethylation of cysteine was set as a fixed modification; serine, threonine and tyrosine phosphorylation and oxidation of methionine were allowed as a variable modification. Mass tolerances for precursor and fragment ions were 10 ppm Da and 0.02 Da, respectively. Abundance ratios were calculated between treatment groups, together with adjusted P-values using the Benjamini-Hochberg method, in Proteome Discoverer.

*Merging of the MiRNA databases*

The miRNA-mRNA interactions were obtained merging 4 public miRNA-mRNA interaction databases (miRDB v6.0^22^, TarBase v9.0^23^, MirTarBase v9.0^24^ and TargetScan v8.0^25^), utilising target genes identified by at least two databases. Nomenclature was aligned to miRBase Release 22^26^.

*Machine learning analysis of miR impact on discordant gene/protein expression*

We divided miRNA-targeted genes into two classes, “class 0”, containing genes whose behaviour is consistent between the transcriptomics and proteomics levels, and “class 1”, where any changes were discordant. More precisely, class 0 comprises (a) genes with no significant changes at either level (mRNA adj p-value > 0.05 and protein adj p-value > 0.05), and (b) genes with changes at both levels in the same direction (mRNA adj p-value < 0.05 and protein adj p-value < 0.05 and mRNAs log2FC > 0 and protein log2FC > 0). Class 1 comprises (a) genes with changes only at the proteomics level (mRNA adj p-value > 0.05 and protein adj p-value ≤ 0.05), (b) genes with changes only at the transcriptomics level (mRNA adj p-value ≤ 0.05 and protein adj p-value > 0.05), and (c) genes with changes at both levels but in opposite directions (mRNA adj p-value < 0.05 and protein adj p-value ≤ 0.05 and mRNAs log2FC > 0 and protein log2FC < 0 or vice versa).

Classes 0 and 1 were defined separately for each biological contrast (OSS vs LSS or ESS vs LSS). Our goal is to develop ML models to predict classes 0 and 1.

- Model features

Four model features were used for the classification. These features are: concentration of the mRNAs and miRNAs (both measured as RPKM counts), net-effect score (calculated as the difference between up-regulated miRNAs and down-regulated miRNAs targeting the same mRNA) and constant of dissociation of each miRNA-mRNA interaction (calculated using Vienna RNA^27^ using the sequences of the miRNAs of the 3’UTR regions of the mRNAs identified using APAtrap^28^)

- Preparation of the dataset

A dataset containing the binary variable and all the model features was prepared for each condition to perform the binary classification. To consider the variability among donors, each gene in each sample was treated as separate entry (individual gene data points). The dataset was then spitted into train set (80%) and test set (20%) using test_train_split function from scikit-learn v1.5^29^. After splitting the data, the train and test set were standardised separately using RobustScaler function from scikit-learn v1.5.

- Training and testing of the model

To optimize model performance, a Random Forest classifier was trained with a 10-fold cross validation and hyperparameter tuning using the RandomForestClassifier and GridSearchCV functions from scikit-learn v1.5. The model was evaluated using the ROC curves, focusing on the AUC score for the overall performance of the model and on the sensitivity to understand the ability of the model to classify class 1 entries.

- Interpretation of the classification

The SHAP values were calculated and visualised using the SHAP library^30^ to interpret the contribution of each feature to the model's predictions.

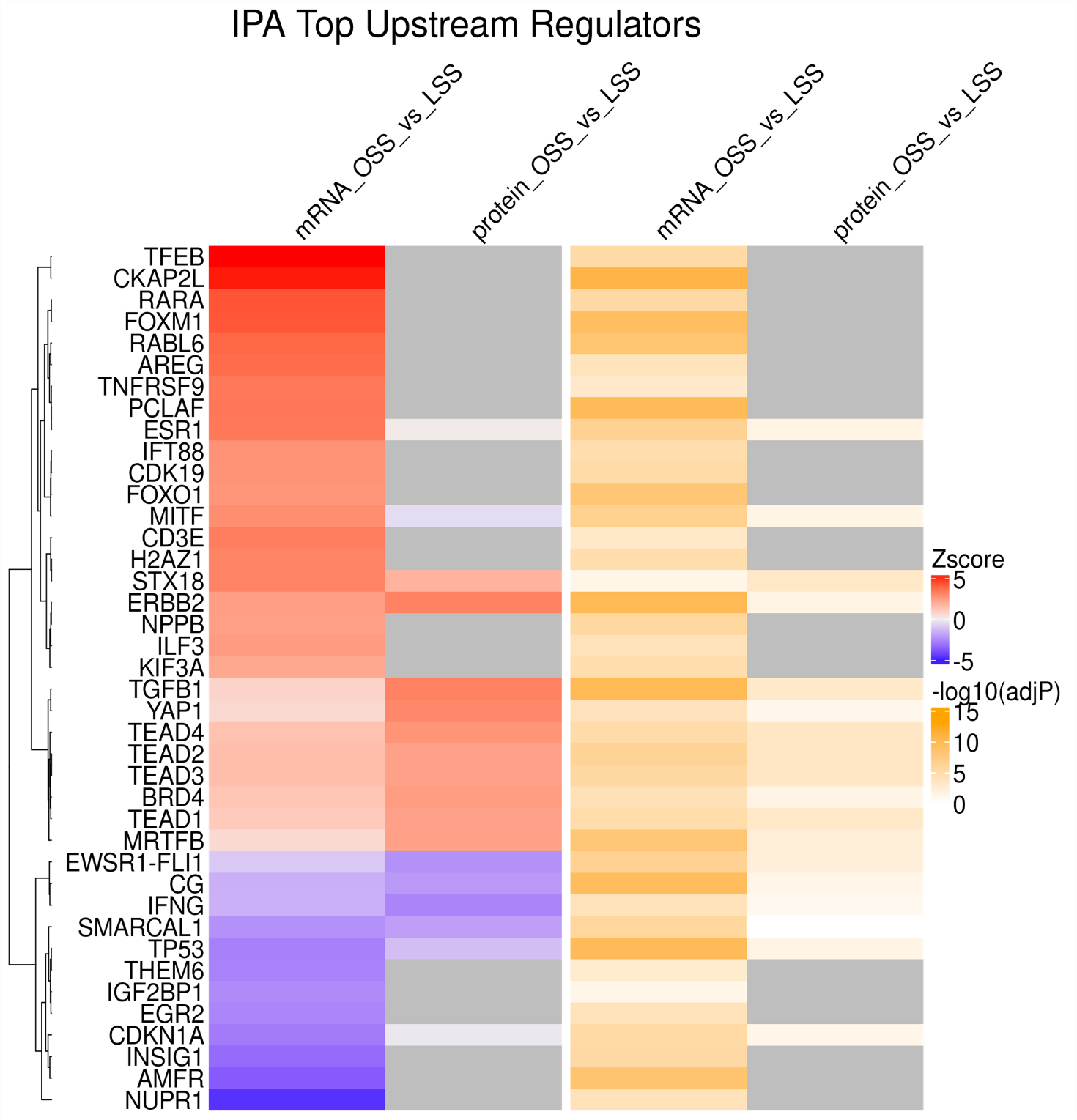

**Figure S1**. Top IPA predicted regulators for genes modulated by OSS compared to LSS.

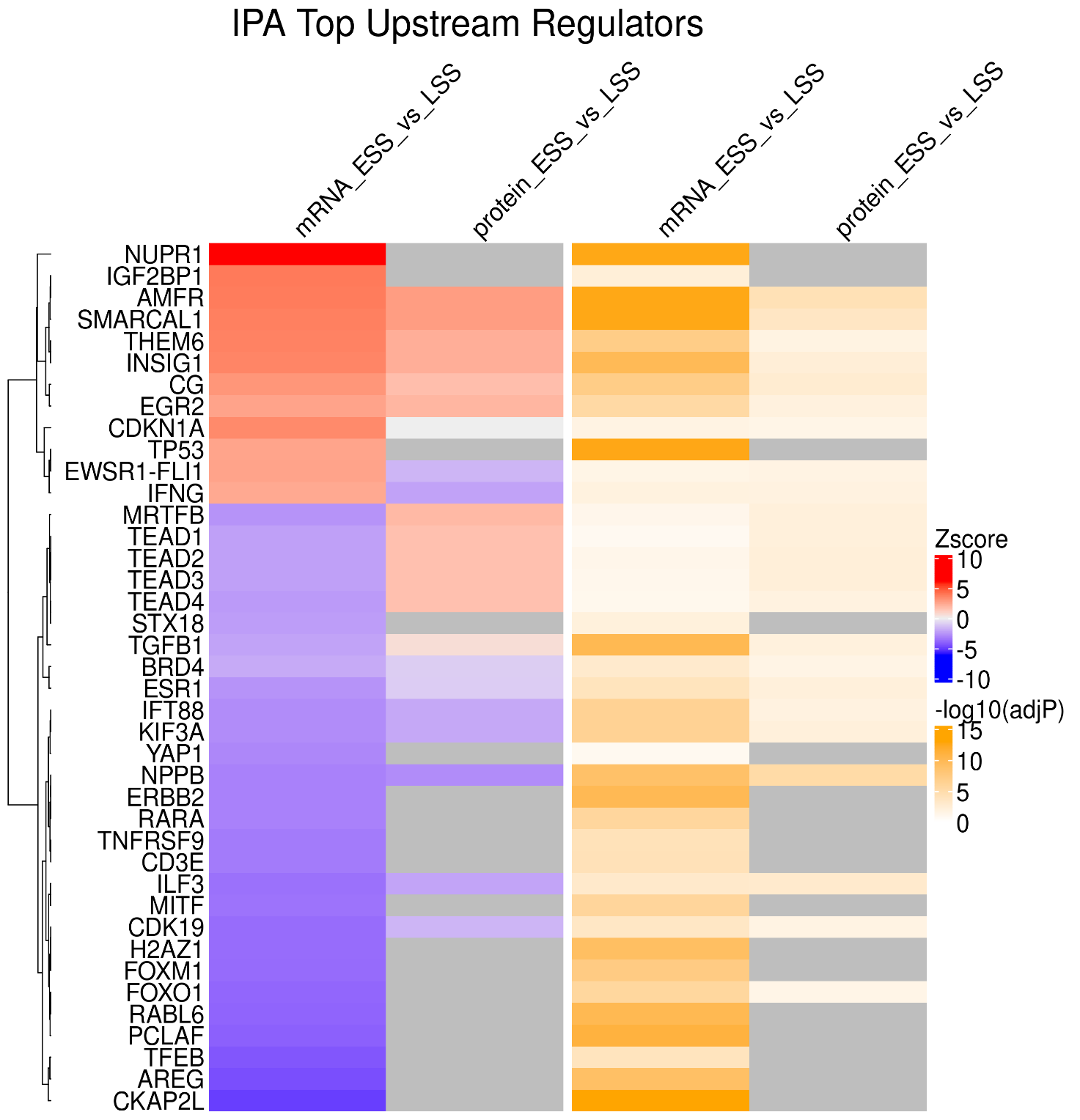

**Figure S2.** Top IPA predicted regulators for genes modulated by OSS compared to LSS.

OSS v LSS

| **ID** | **PATHWAY NAME** |
| --- | --- |
| 1 | Cholesterol biosynthesis |
| 2 | Superpathway of Cholesterol Biosynthesis |
| 3 | Cholesterol Biosynthesis I |
| 4 | Cholesterol Biosynthesis II (via 24,25-dihydrolanosterol) |
| 5 | Cholesterol Biosynthesis III (via Desmosterol) |
| 6 | Pathogen Induced Cytokine Storm Signaling Pathway |
| 7 | Kinetochore Metaphase Signaling Pathway |
| 8 | Activation of gene expression by SREBF (SREBP) |
| 9 | Tumor Microenvironment Pathway |
| 10 | Mitotic Roles of Polo-Like Kinase |
| 11 | Keratinization |
| 12 | Pulmonary Fibrosis Idiopathic Signaling Pathway |
| 13 | Class A/1 (Rhodopsin-like receptors) |
| 14 | Airway Pathology in Chronic Obstructive Pulmonary Disease |
| 15 | Mevalonate Pathway I |
| 16 | Superpathway of Geranylgeranyldiphosphate Biosynthesis I (via Mevalonate) |
| 17 | Mitotic Prometaphase |
| 18 | Potassium Channels |
| 19 | LXR/RXR Activation |
| 20 | Agranulocyte Adhesion and Diapedesis |

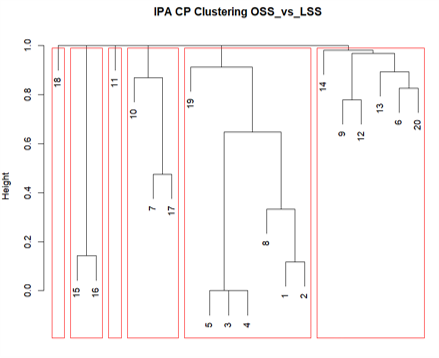

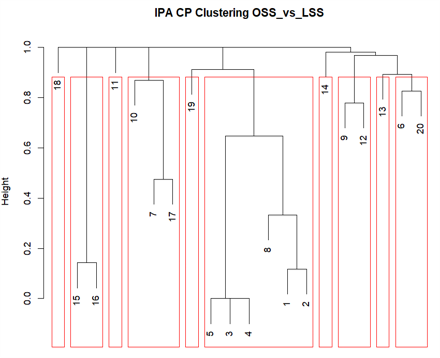

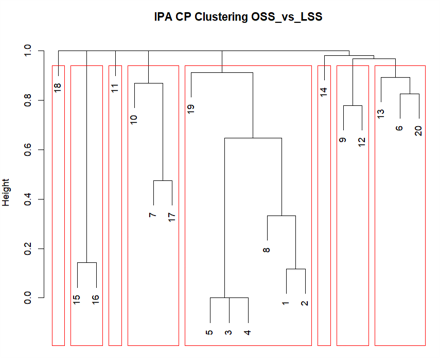

**A**

**C**

**B**

**Figure S3 Dendrograms of pathway clustering on the OSS vs LSS condition –** generated in R using binary distance between the gene sets associated with each pathway, and hierarchical clustering via the hclust algorithm. Red rectangles indicate the division into: **(A)** 6 clusters, **(B)** 8 clusters and **(C)** 10 clusters.

**A**

| **ID** | **PATHWAY NAME** |
| --- | --- |
| 1 | Cholesterol biosynthesis |
| 2 | Superpathway of Cholesterol Biosynthesis |
| 3 | Cholesterol Biosynthesis I |
| 4 | Cholesterol Biosynthesis II (via 24,25-dihydrolanosterol) |
| 5 | Cholesterol Biosynthesis III (via Desmosterol) |
| 6 | Pathogen Induced Cytokine Storm Signaling Pathway |
| 7 | Kinetochore Metaphase Signaling Pathway |
| 8 | Activation of gene expression by SREBF (SREBP) |
| 9 | Tumor Microenvironment Pathway |
| 10 | Mitotic Roles of Polo-Like Kinase |
| 11 | Keratinization |
| 12 | Pulmonary Fibrosis Idiopathic Signaling Pathway |
| 13 | Class A/1 (Rhodopsin-like receptors) |
| 14 | Airway Pathology in Chronic Obstructive Pulmonary Disease |
| 15 | Mevalonate Pathway I |
| 16 | Superpathway of Geranylgeranyldiphosphate Biosynthesis I (via Mevalonate) |
| 17 | Mitotic Prometaphase |
| 18 | Potassium Channels |
| 19 | LXR/RXR Activation |
| 20 | Agranulocyte Adhesion and Diapedesis |

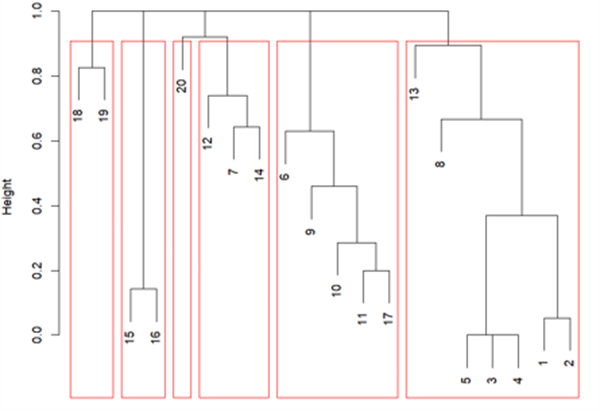

**B**

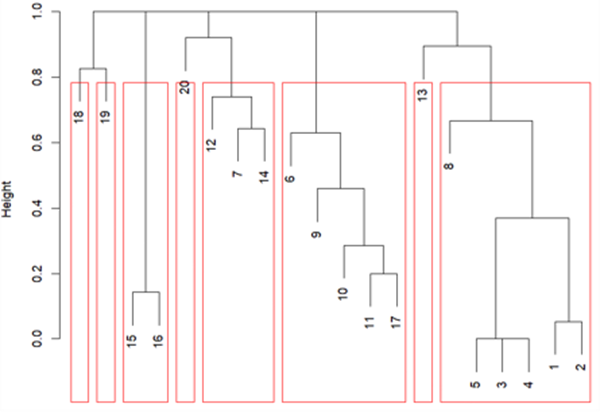

**C**

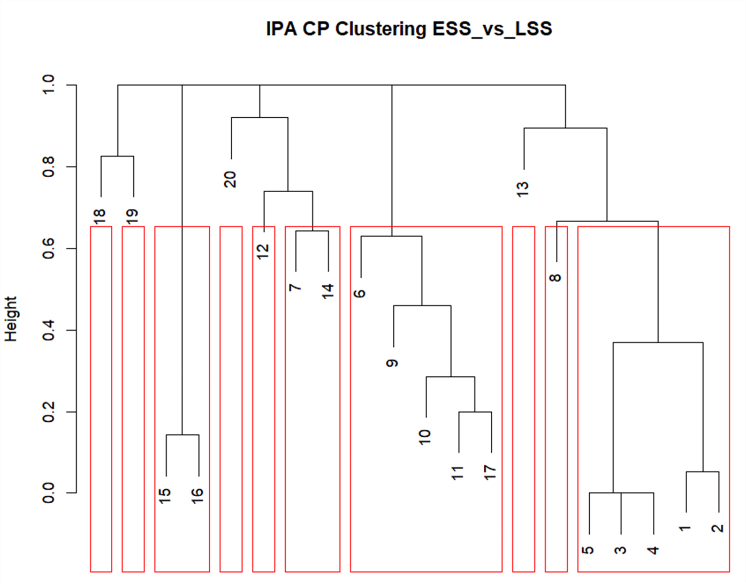

**Figure S4 Dendrograms of pathway clustering on the ESS vs LSS condition –** generated in R using binary distance between the gene sets associated with each pathway, and hierarchical clustering via the hclust algorithm. Red rectangles indicate the division into: **(A)** 6 clusters, **(B)** 8 clusters and **(C)** 10 clusters.

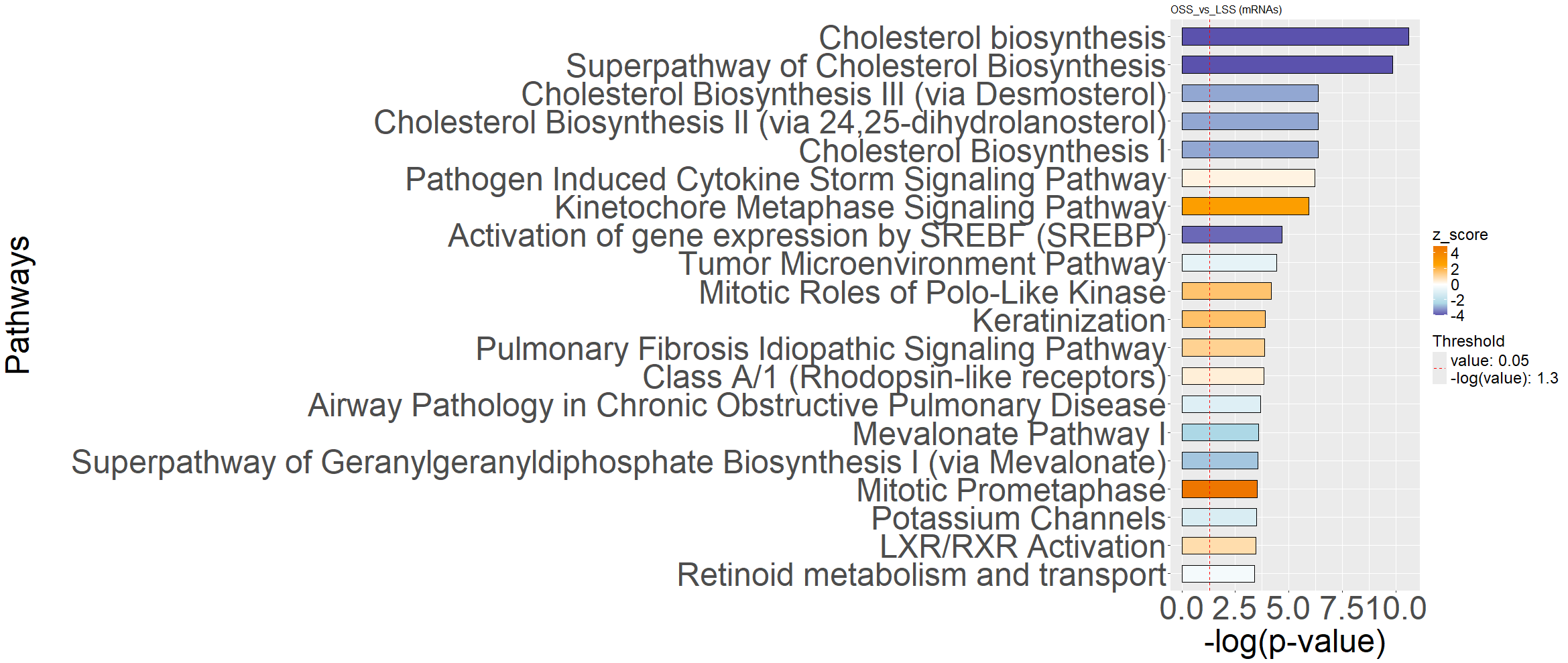

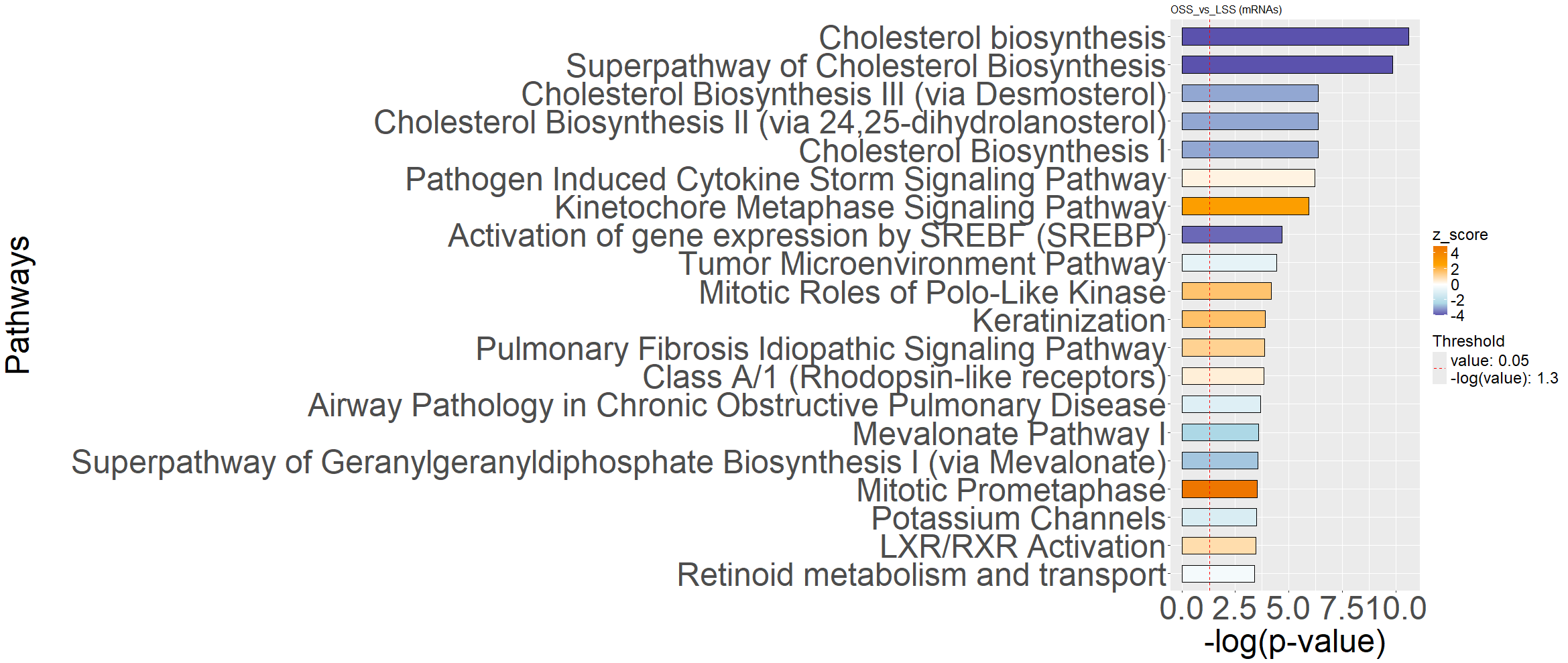

**Figure S5.** Uncondensed pathway analysis of genes with altered expression in oscillatory flow. Generated by IPA

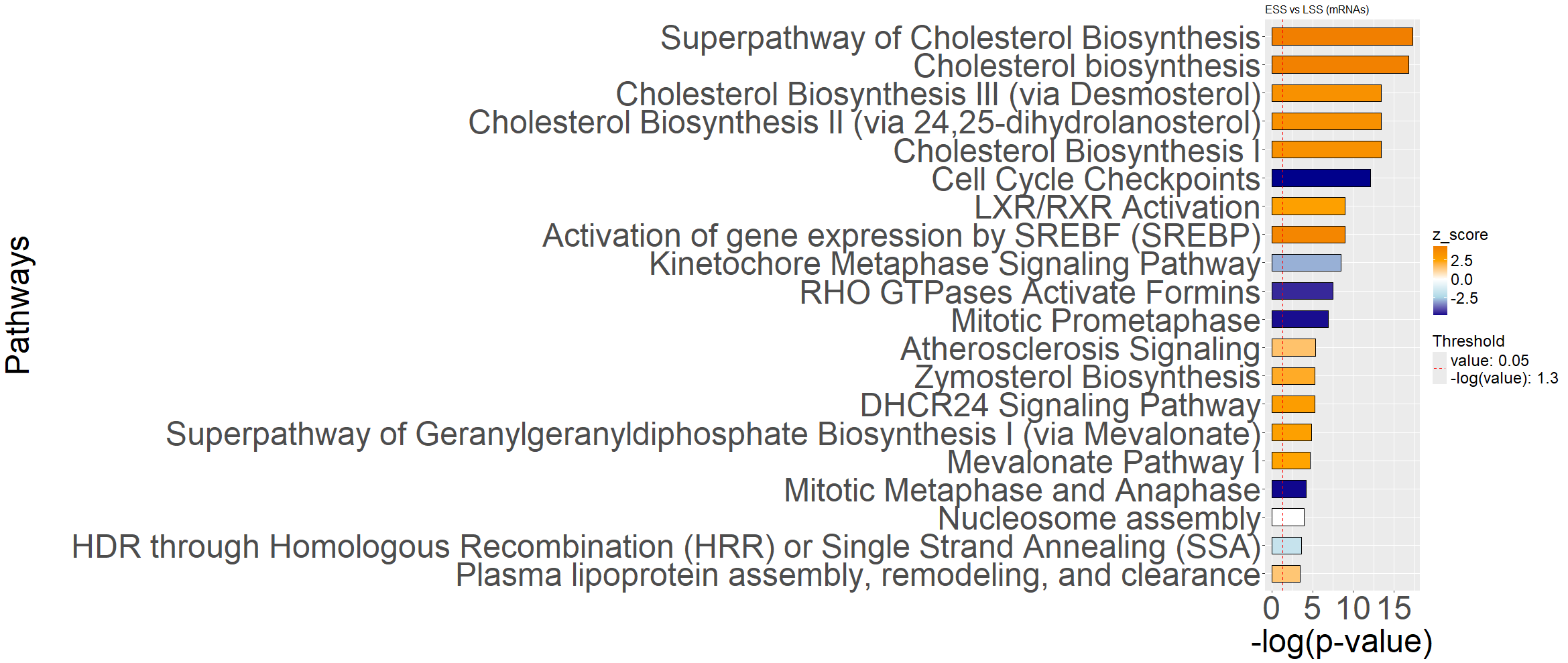

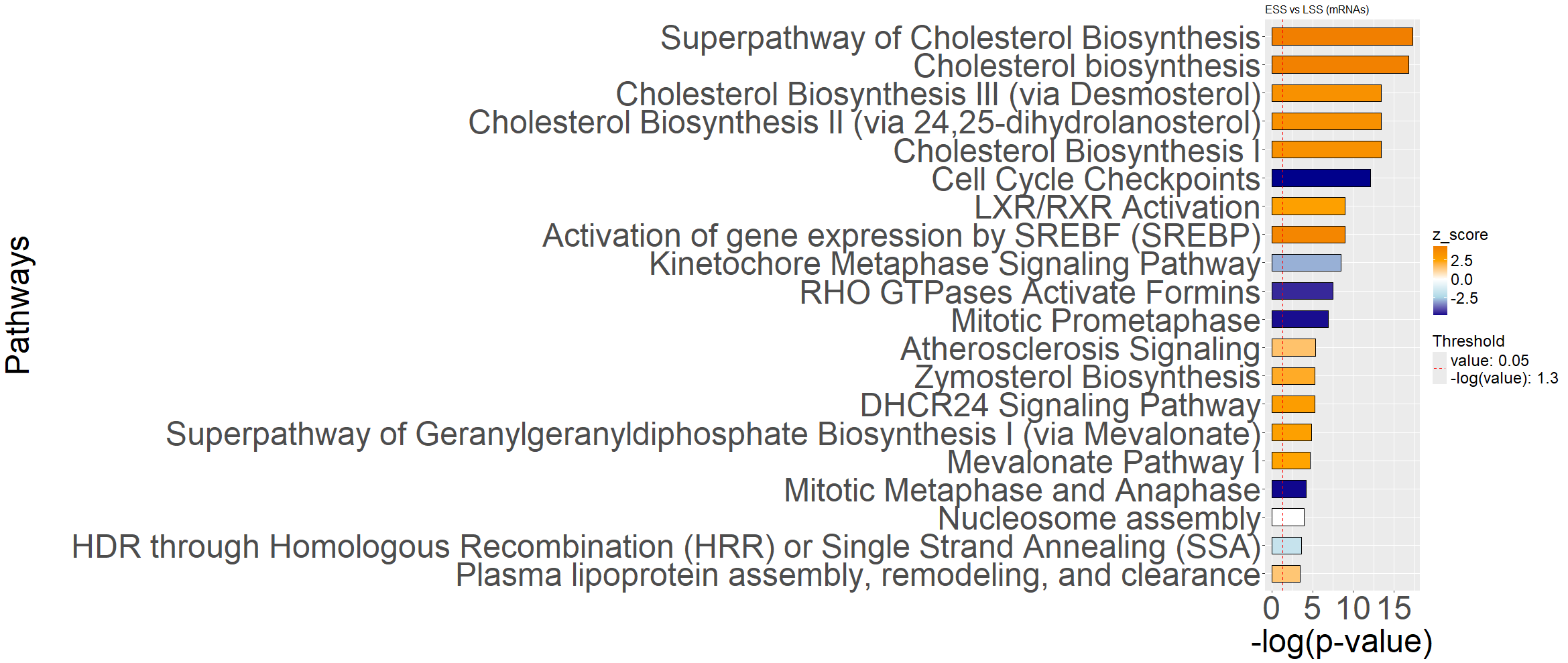

**Figure S6.** Uncondensed pathway analysis of genes with altered expression in elevated flow. Generated by IPA

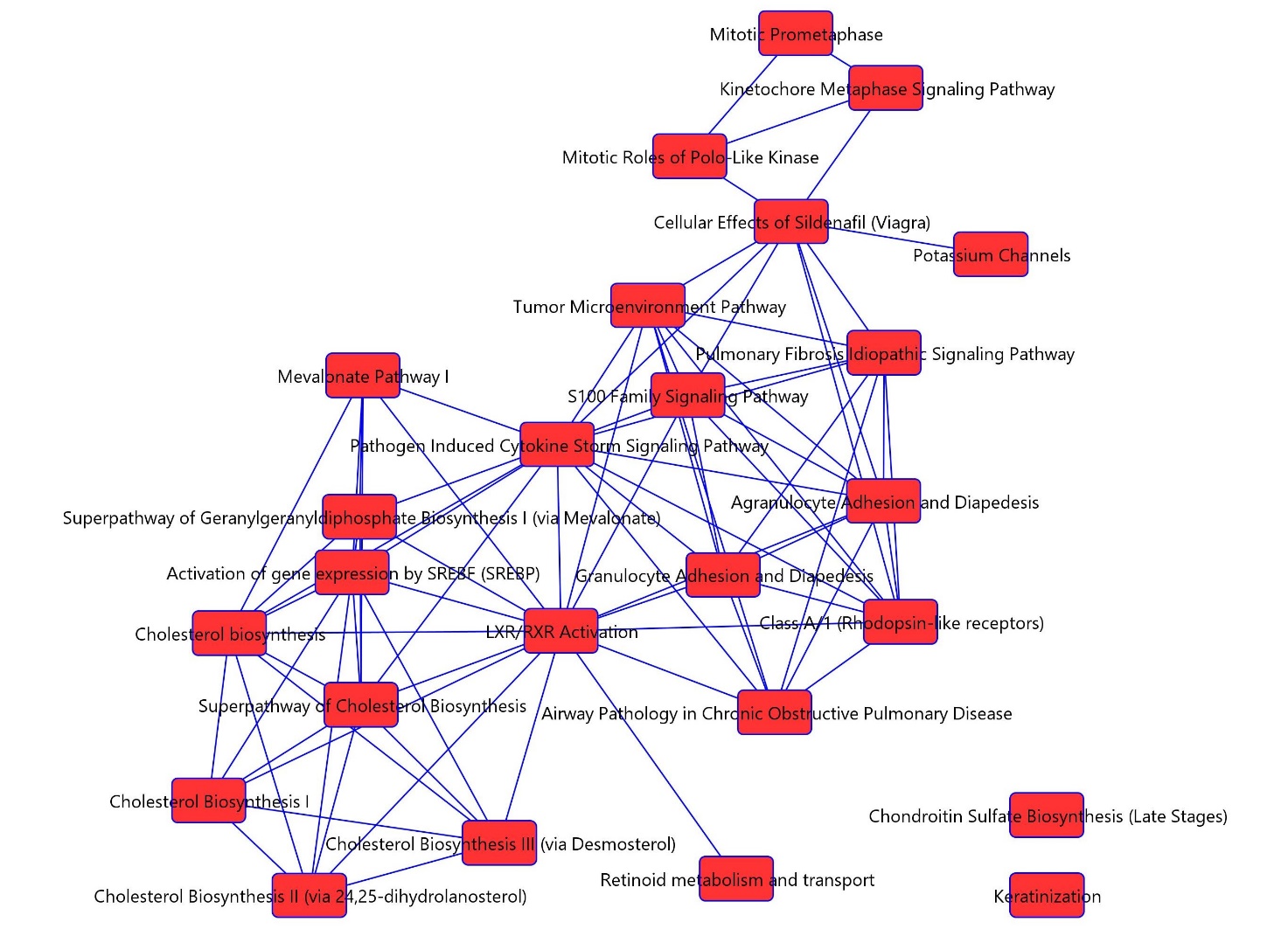

**Figure S7.** Networks of overlapping IPA canonical pathways, OSS.

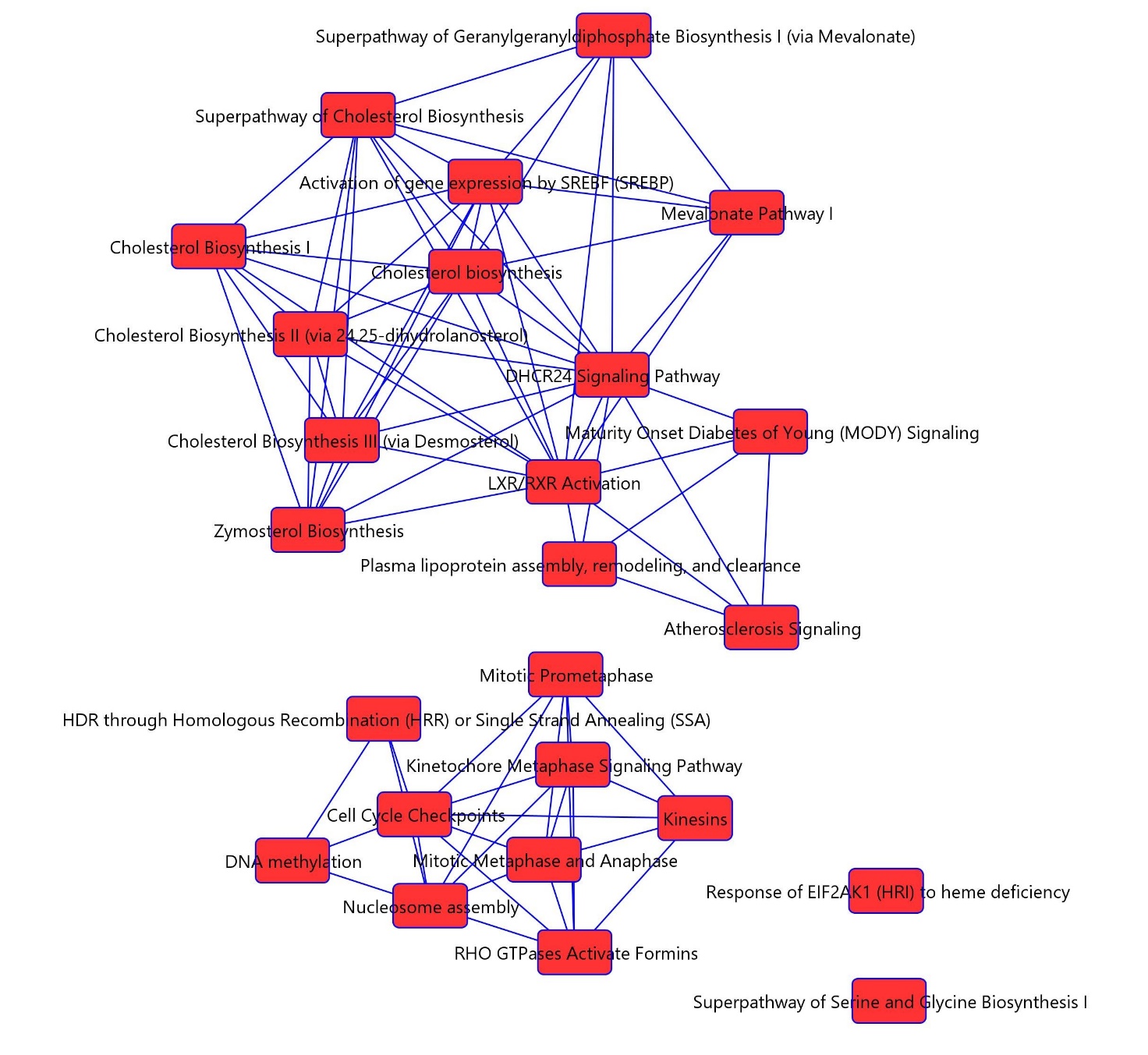

Figure S8. Networks of overlapping IPA canonical pathways, B: ESS.

**OSS vs LSS**

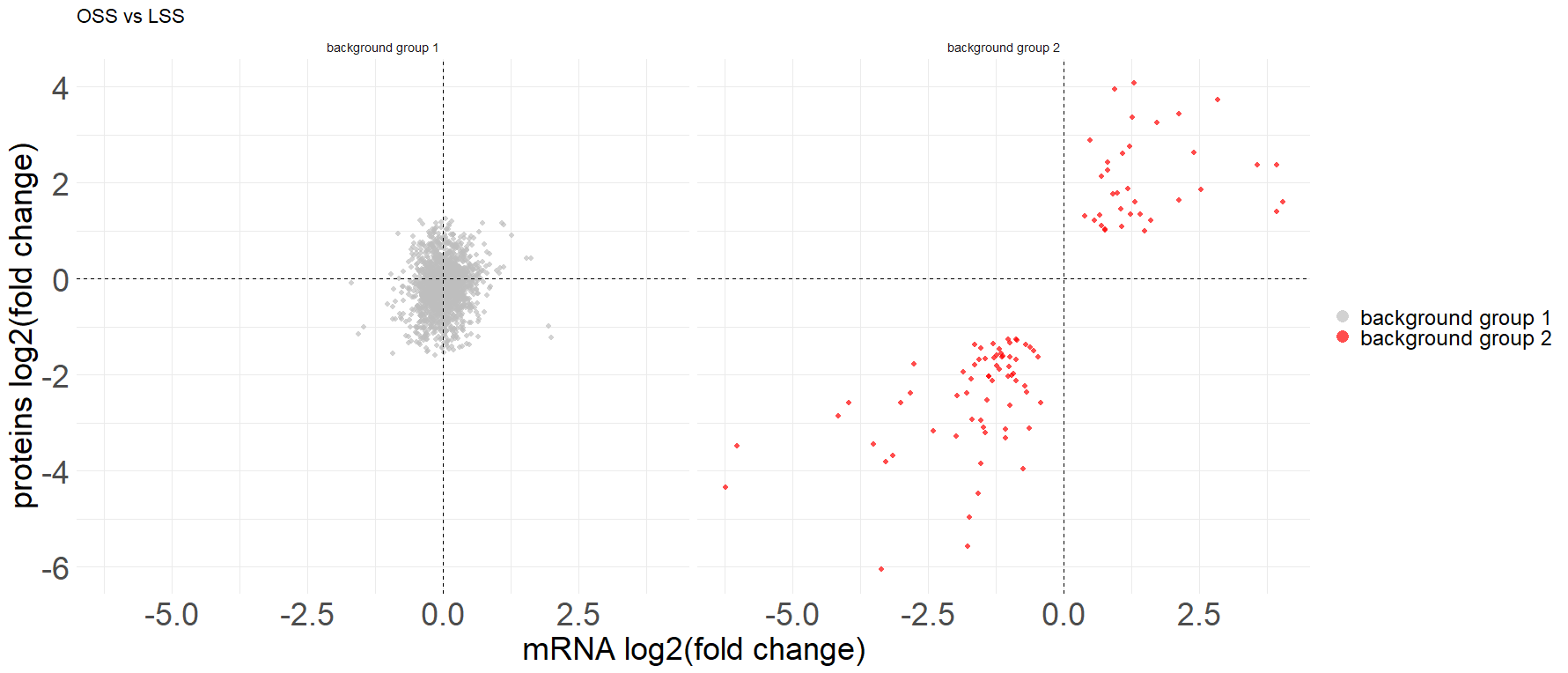

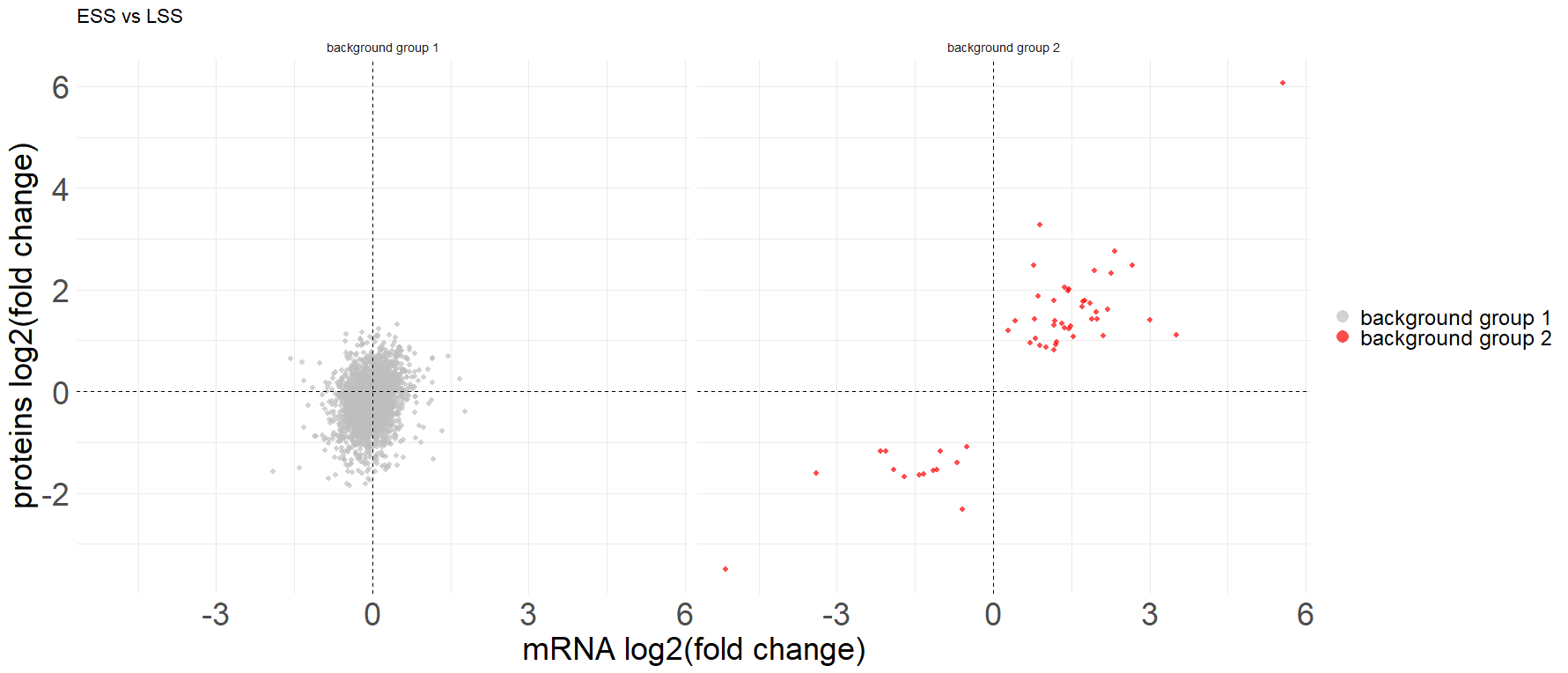

**ESS vs LSS**

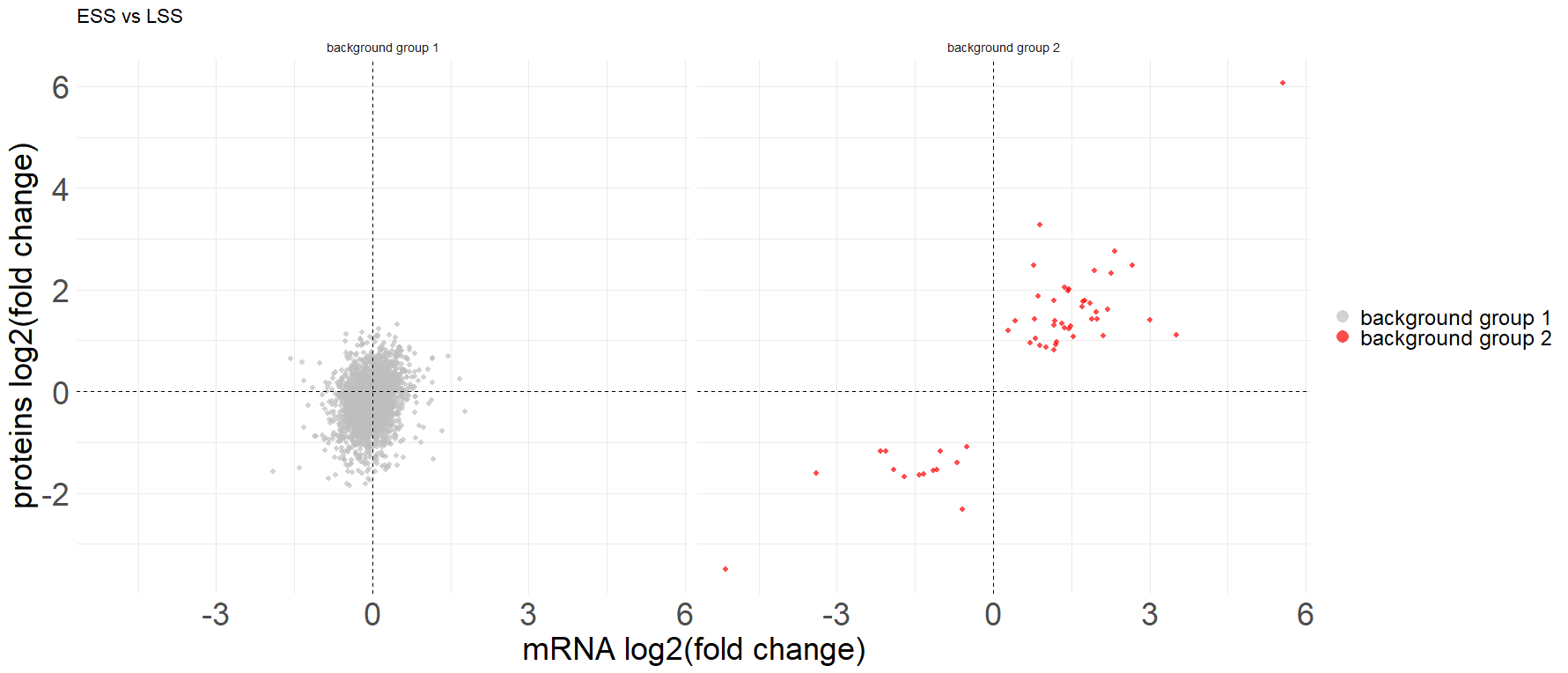

**Figure S9. Scatter plots of the background groups.** The plots show the relationship between the log2FC of the genes (x-axis) and the log2FC of the associated proteins (y-axis). Each gene and the respective protein are represented by a single dot. The colour of the dots represents background group 1 (grey) and background group 2 (red). At the top, genes in background group 1 and 2 are defined for OSS vs LSS. Lower figures, genes in background group 1 and 2 are defined for ESS vs LSS.

**OSS vs LSS	ESS vs LSS**

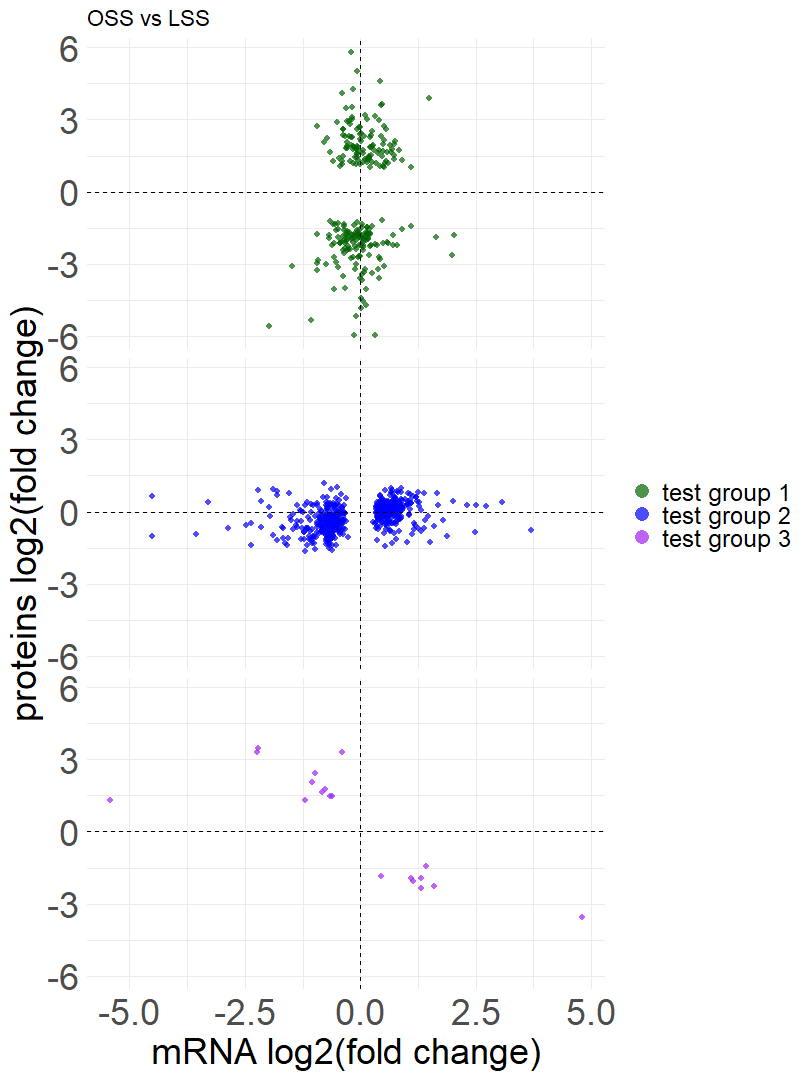

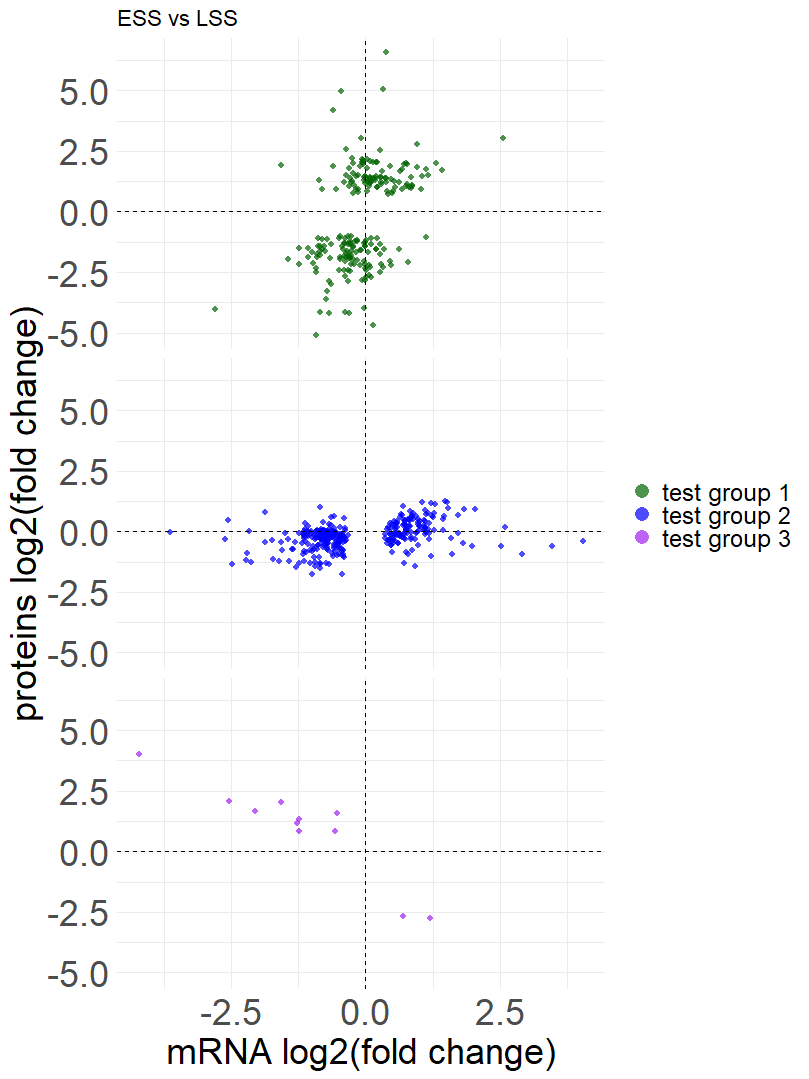

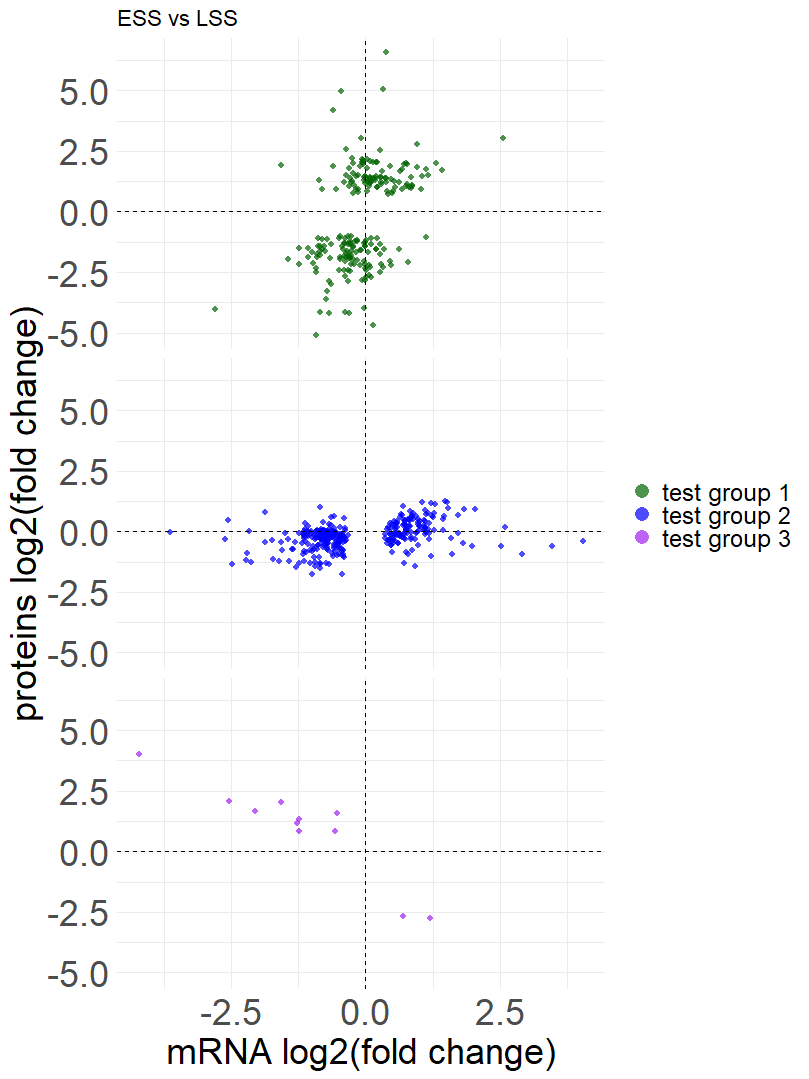

**Figure S10. Scatter plots of the test groups.** The plots show the relationship between the log2FC of the genes (x-axis) and the log2FC of the associated proteins (y-axis). Each gene and the respective protein are represented by a single dot. The color of the dots represents test group 1 (green), test group 2 (blue) and test group 3 (purple). On the left, genes in test group 1, 2 and 3 are defined based on the OSS vs LSS contrast. On the right, genes in test group 1, 2 and 3 are defined based on the ESS vs LSS contrast.

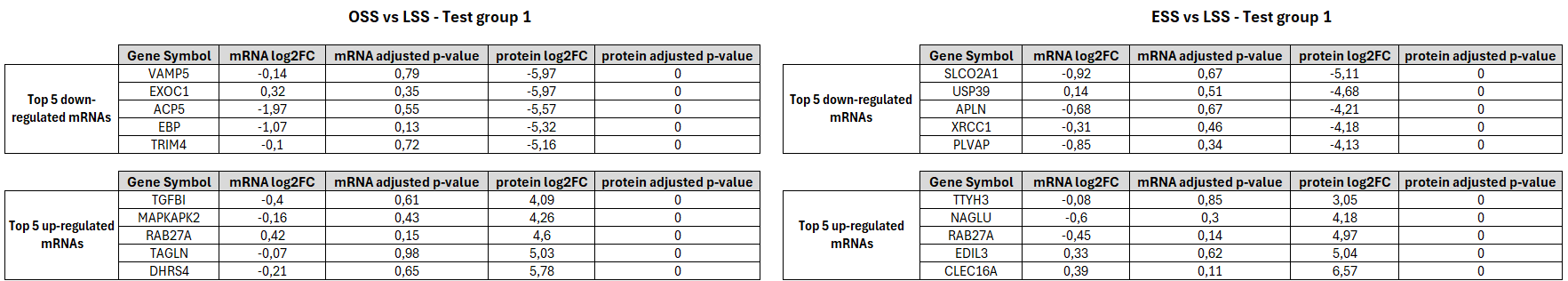

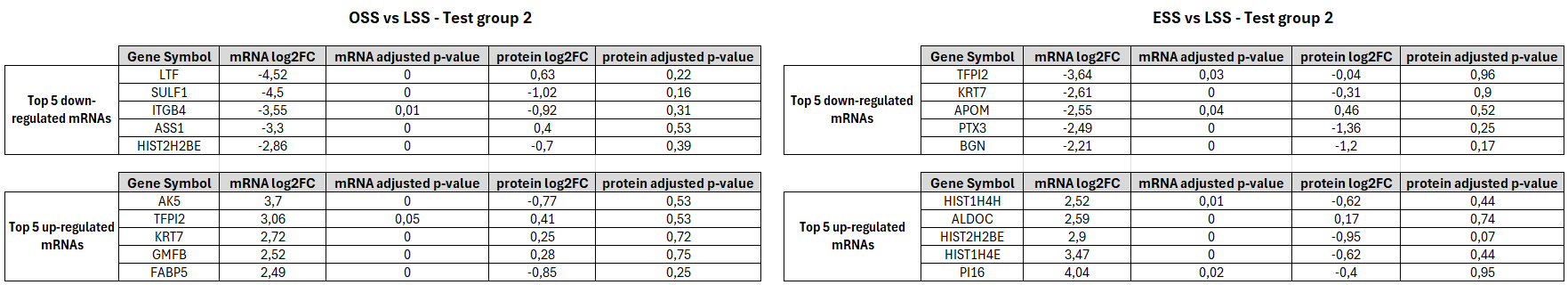

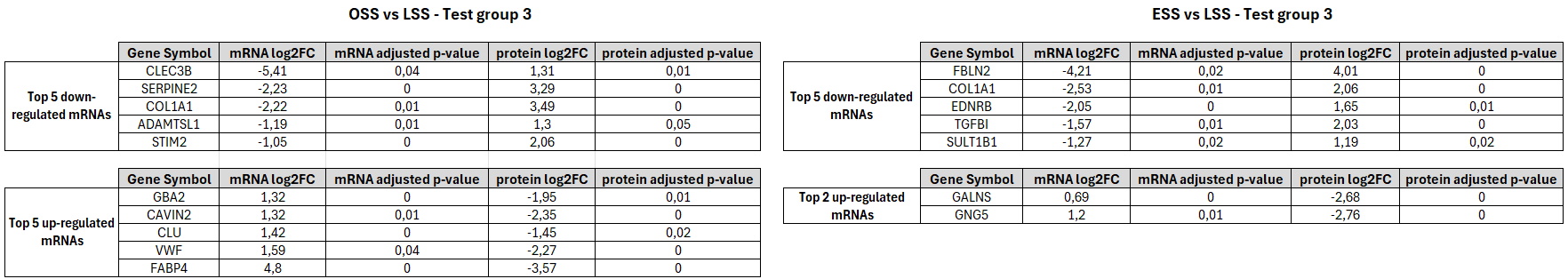

**Table S1.** Top 5 up- and down-regulated genes in each discordantly regulated gene test group for both OSS and ESS conditions**.**

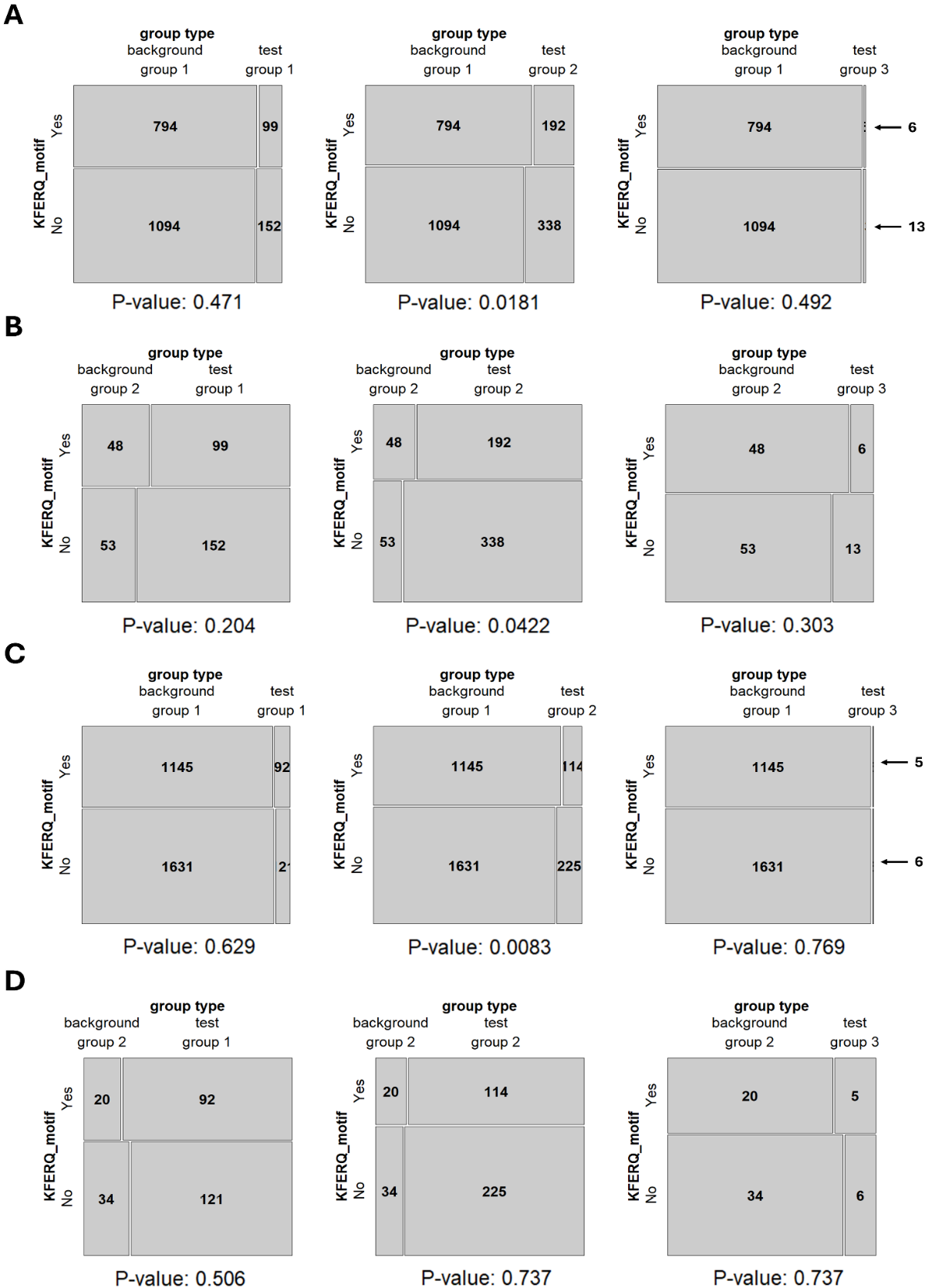

**Figure S11. Tests of Association for KFERQ-like motifs**

To investigate the relationship between miRNA impact and discordant changes in the previously defined test groups, we assessed associations using either Fisher’s exact test or the Chi-square test, depending on the sample size. None of the tests were found to be significant, as all were associated with a p-value greater than 0.05. As a result, there was no evidence of a significant association between a gene belonging to one of the discordantly regulated expression groups and its predicted inclusion of a KFERQ motif

**OSS vs LSS**

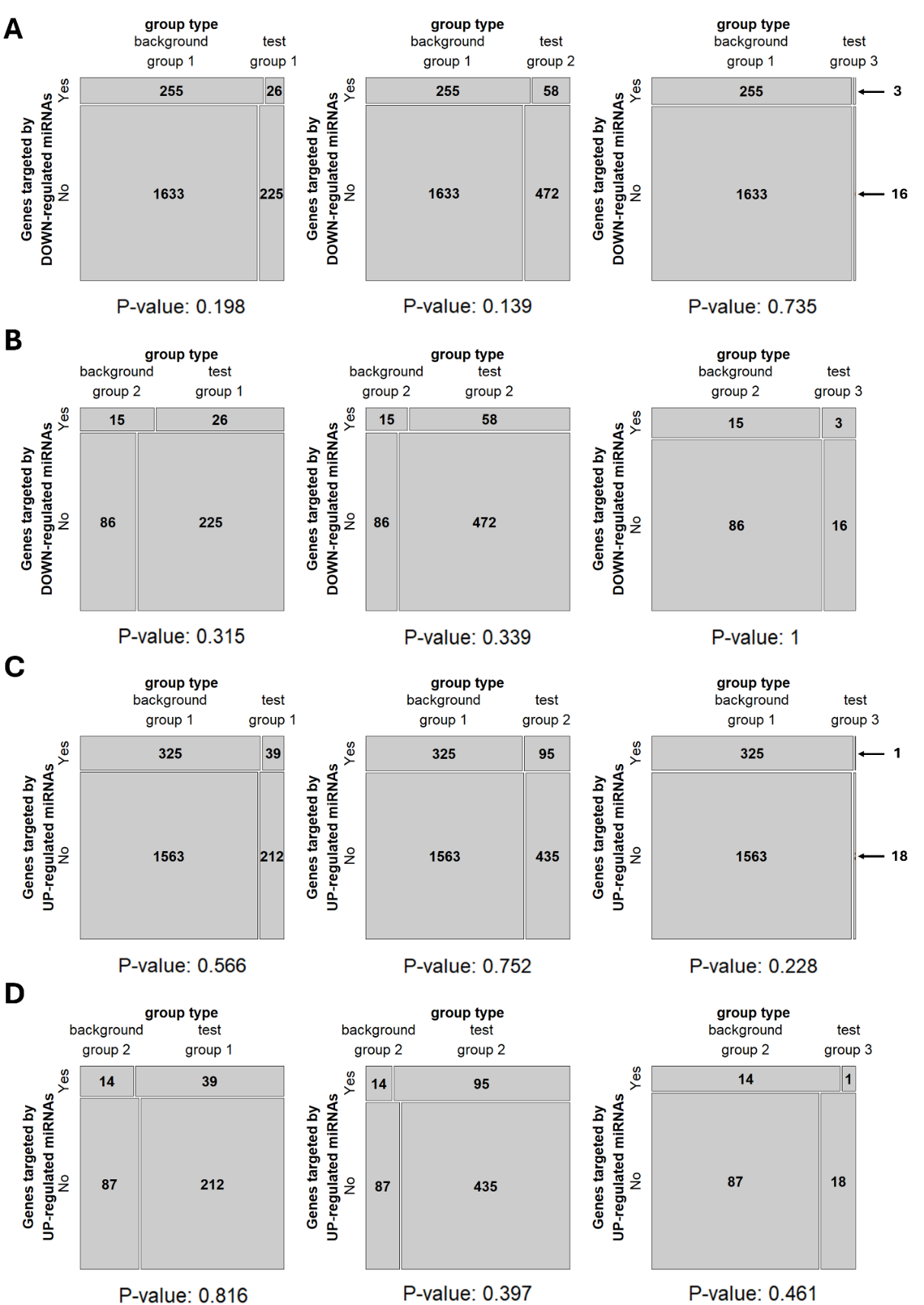

**ESS vs LSS**

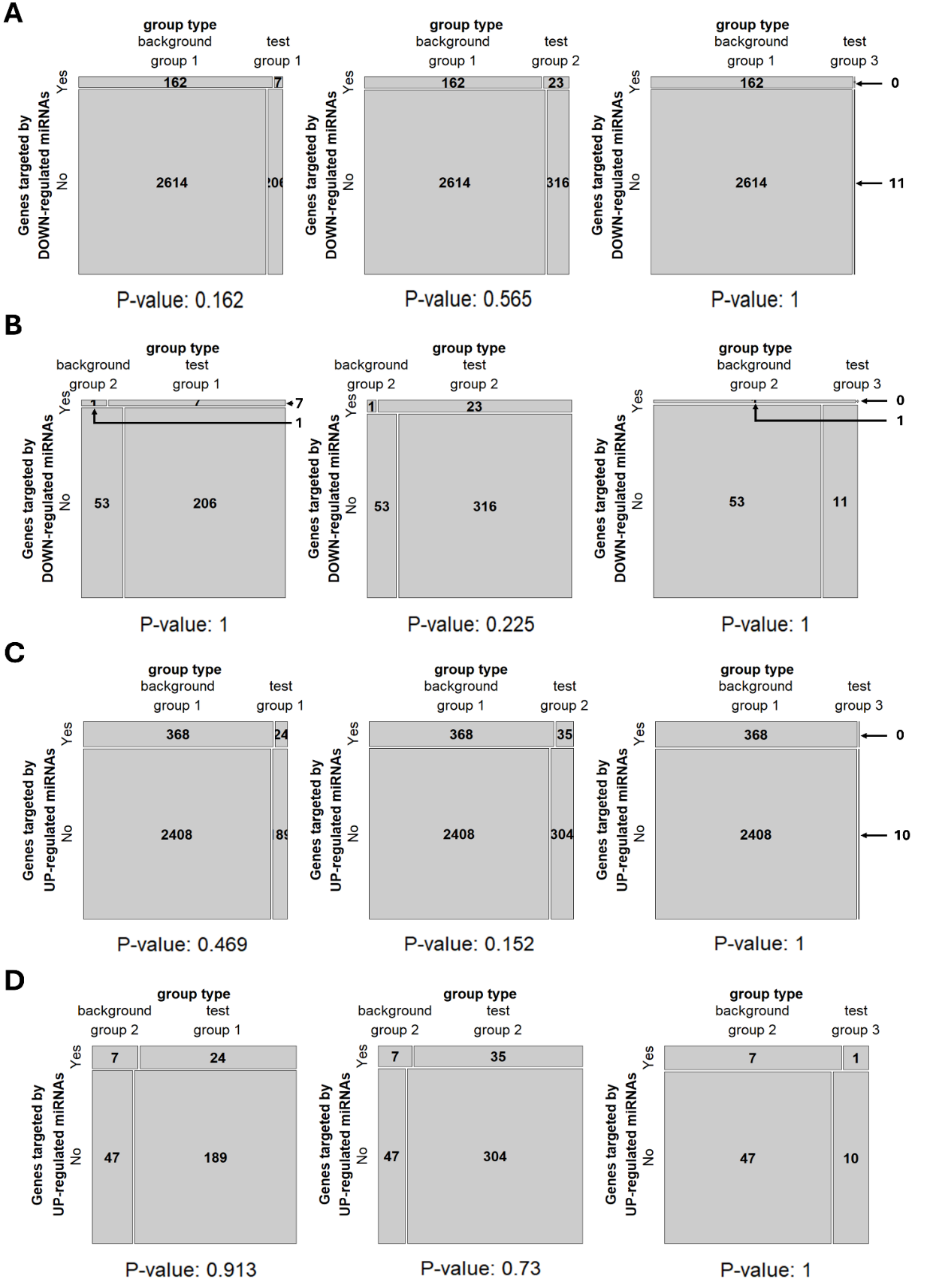

**Figure S12. Tests of Association for predicted miR targets**

To investigate the relationship between miRNA impact and discordant changes in the previously defined test groups, we assessed associations using either Fisher’s exact test or the Chi-square test, depending on the sample size. None of the tests were found to be significant, as all were associated with a p-value greater than 0.05. As a result, there was no evidence of a significant association between a gene belonging to one of the discordantly regulated test groups and its predicted targeting by an up-regulated or down-regulated miRNA.
